# Atypical B cells are a normal component of immune responses to vaccination and infection in humans

**DOI:** 10.1101/2020.05.28.120808

**Authors:** Henry J. Sutton, Racheal Aye, Azza H. Idris, Rachel Vistein, Eunice Nduati, Oscar Kai, Jedida Mwacharo, Xi Li, Xin Gao, T. Daniel Andrews, Marios Koutsakos, Thi H. O. Nguyen, Maxim Nekrasov, Peter Milburn, Auda Ethala, Andrea A. Berry, KC Natasha, Sumana Chakravarty, B. Kim Lee Sim, Adam K. Wheatley, Stephen J. Kent, Stephen L. Hoffman, Kirsten E. Lyke, Philip Bejon, Fabio Luciani, Katherine Kedzierska, Robert A. Seder, Francis M. Ndungu, Ian A. Cockburn

## Abstract

The full diversity of the circulating human B cell compartment is unknown. Flow cytometry analysis suggests that in addition to naïve and memory B cells, there exists a population of CD11c^+^, CD27^−^ CD21^−^ “atypical” B cells, that are associated with chronic or recurrent infection and autoimmunity. We used single cell RNA-seq approaches to examine the diversity of both antigen-specific B cells and total B cells in healthy subjects and individuals naturally-exposed to recurrent malaria infections. This analysis revealed two B cell lineages: a classical lineage of activated and resting memory B cells, and an atypical-like lineage. Surprisingly, the atypical lineage was common in both malaria exposed individuals and non-exposed healthy controls. Using barcoded antibodies in conjunction with our transcriptomic data, we found that atypical lineage cells in healthy individuals lack many atypical B markers and thus represent an undercounted cryptic population. We further determined using antigen specific probes that atypical cells can be induced by primary vaccination in humans and can be recalled upon boosting. Collectively these data suggest that atypical cells are not necessarily pathogenic but can be a normal component of B responses to antigen.

## Introduction

The majority of currently approved vaccines require the generation of an effective antibody response to provide long term immune protection (Plotkin, 2010). An effective antibody response requires the formation of germinal centers (GCs) to produce somatically hypermutated and affinity matured long-lived plasma cells (LLPCs) that secrete high-affinity antibody as well as antigen-experienced “memory” B cells (MBCs) that are primed to produce a faster, larger and more effective response upon secondary exposure (Tangye et al., 2003). In humans, circulating human B cells have been classified based on the expression of the surface proteins CD38, CD27 and CD21. Plasma cells (PCs) express high levels of CD38 and CD27 (Horst et al., 2002) while CD27^+^ CD21^+^ B cells are considered to be MBCs (Klein et al., 1997; Tangye et al., 1998). These CD27^+^ cells show high levels of affinity maturation and readily differentiate into antibody secreting PCs after stimulation compared to CD27^−^ CD21^+^ naïve cells (Good et al., 2009; Tangye et al., 2003). Populations of CD27^+^, CD21^−^ B cells have also been described, which have previously been associated with an activated B cell phenotype, predisposed to differentiate into PCs (Avery et al., 2005; Lau et al., 2017). Conversely a subset of B cells that are CD27^−^ CD21^−^ have also been observed, originally in tonsils and later in peripheral blood (Ehrhardt et al., 2005; Fecteau et al., 2006). These cells, commonly referred to as atypical B cells (atBC), are observed at high frequencies in conditions of chronic antigen stimulation such as infection with HIV or malaria (Moir et al., 2008; Weiss et al., 2009) or autoimmune disease (Isnardi et al., 2010; Wei et al., 2007).

Because they are often found in chronic infection and autoimmune disease, atBCs are usually considered to be a population of anergic or exhausted B cells that arise due to chronic antigenic stimulation. In support of this, atBCs typically express high levels of inhibitory receptors such as those belonging to the family of Fc-receptor-like (FCRL) molecules, as well as having muted BCR signaling and display little to no capacity to differentiate into PCs following BCR stimulation *in vitro* (Moir et al., 2008; Portugal et al., 2015; Sullivan et al., 2015). However, not all data indicate that atBCs are an exhausted population. While SLE patients with high disease scores carry high numbers of atBCs, a recent study suggested that these are short-lived activated cells, in the process of differentiating into PCs (Jenks et al., 2018). Similarly, it has been shown that BCRs used by atBCs specific to *Plasmodium falciparum* could also be found contributing to the anti-*P. falciparum* antibody response (Muellenbeck et al., 2013). Furthermore, studies in mice show that short-lived, recently activated B cells express high levels of CD11c^+^ and have similar gene expression patterns to human atBCs (Kim et al., 2019; Perez-Mazliah et al., 2018).

To better understand the heterogeneity of the circulating B cell response in humans, and gain insight into the role of atBCs, we performed single cell RNA-seq on antigen-specific B cells from malaria exposed adults, we compared these data to single-cell RNA-seq on non-antigen specific B cells from both malaria-exposed and non-exposed individuals. Finally, we examined the phenotypic profile when antigen specific atBCs and MBCs arise in the response to vaccination. Collectively we found that even non-exposed individuals carry high numbers of cells that express an atBC transcriptomic signature. Our vaccination studies revealed that antigen-specific atBCs arise following a primary immune response and are able to respond upon secondary exposure, suggesting that atBCs may have functional role in the human humoral response.

## Results

### Single cell RNA-seq reveals three distinct populations of antigen-experienced circulating B cells

The initial studies focused on the transcriptional diversity of the circulating B cell populations in malaria-vaccinated and -exposed humans by single-cell RNA sequencing (scRNA-seq). Specifically, we isolated CD19^+^, CD20^+^ B cells that (i) were class switched i.e. IgD^−^ and (ii) bound specific antigens. *P. falciparum* circumsporozoite (PfCSP) specific cells were isolated from the peripheral blood of five Kenyan children 6.5 and 74 months after receiving the last dose of the CSP-based RTS,S vaccine. To further examine the response to natural exposure to malaria PfCSP specific B cells, as well as B cells specific for the *P. falciparum* merozoite surface protein-1 (PfMSP1) were also sorted from 6 adult Kenyans from an area of moderate to high malaria transmission (Figure 1A); details of study subjects are given in Table S1). Individuals in this population carry high numbers of circulating atBCs as described using the absence of CD21 and CD27 as markers (Aye et al., 2020). We also sorted tetanus toxoid (TT) specific B cells from the adult subjects; which we have previously shown to have a more classical MBC phenotype (Aye et al., 2020). Because B cells specific for a given antigen are rare within an individual, only ∼10-50 antigen specific cells could be sorted per sample. Accordingly, we used a modified version of the relatively low throughput Smart-seq2 protocol (Picelli et al., 2014) to obtain transcriptomes of the individual cells.

**Figure 1:**
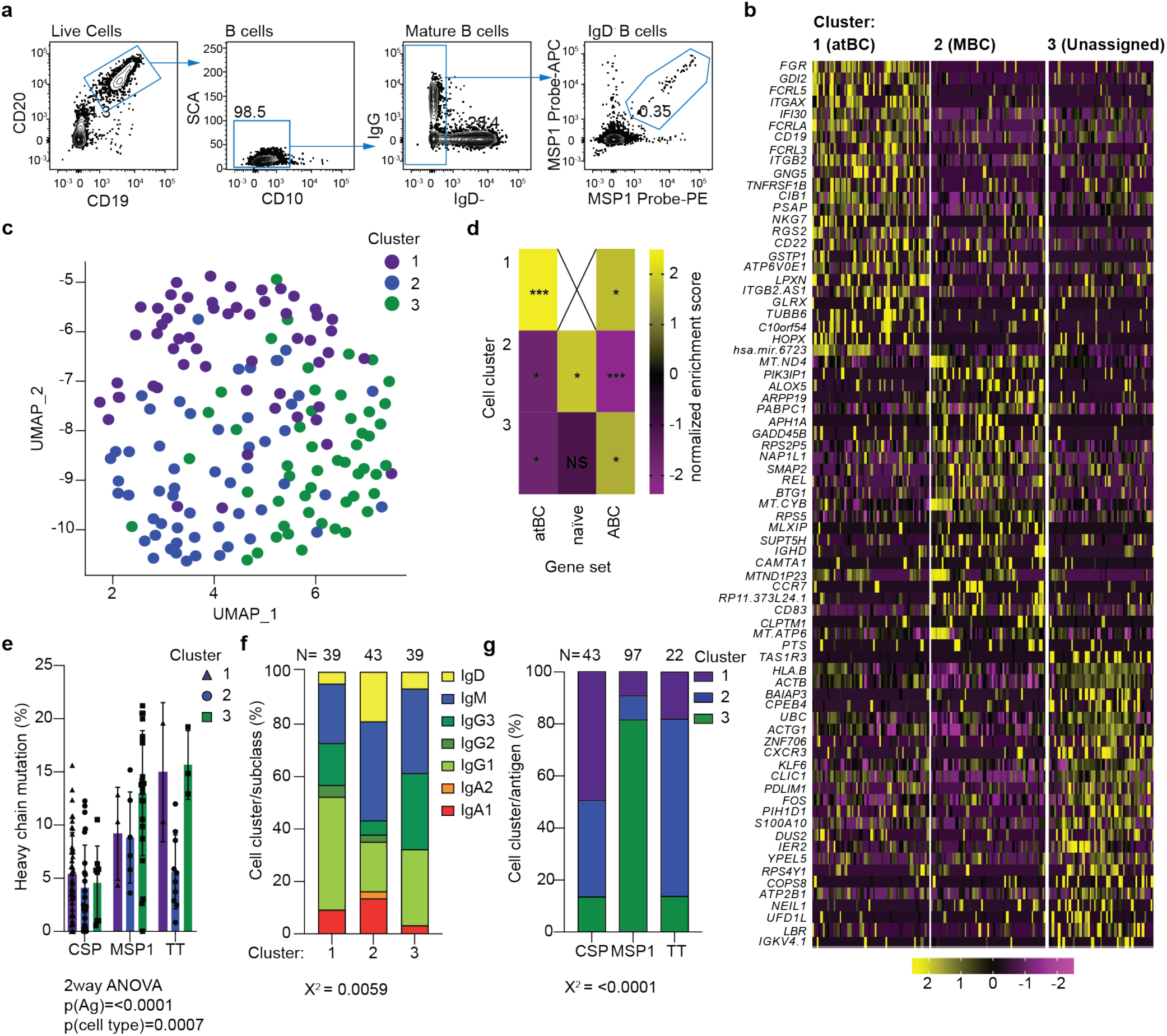
Three distinct populations of antigen-experienced B cells revealed by single cell RNA-seq. CSP, MSP1 and TT specific B cells were index, single cell sorted from malaria vaccinated and exposed donors; transcriptomic information for each cell was generated using Smart-seq2 methodology. **A**. Representative flow cytometry plots showing the gating strategy used to sort mature IgD-antigen-specific B cells. **B**. Heatmap showing the expression of the top 25 DEGs (row) per cluster for each cell (column). **C**. Unsupervised clustering of circulating antigen-specific B cells visualized using UMAP. Each point represents a cell and is colored by cluster. **D**. Heatmap displaying the normalized enrichment scores of multiple GSEA comparing each cluster vs previously published gene sets from atBCs, naïve B cells and ABCs **E**. Percentage of antigen-specific cells that were found in each cluster, analysis was by chi-squared test on the absolute values which are given above each bar. **F**. Percentage of antibody isotype usage by each cluster, analysis was by chi-squared test on the absolute values which are given above each bar. **G**. Percentage of mutations found in the heavy chain V(D)J region of each antigen-specific cell per cluster, analysis was by 2-way ANOVA including each subject as a blocking factor, bars represent mean ± s.d.. Where the exact p value is not quoted * p<0.05, **p<0.01, ***P<0.001.

Following quality control steps, a total of 163 transcriptomes from the 11 individuals were obtained and pooled for analysis using the R package, *Seurat* (Butler et al., 2018). Unsupervised hierarchical clustering grouped the cells into 3 clusters (Figure 1B-C). Differentially expressed genes (DEGs) were identified for each cluster using the Wilcox test to calculate the difference between the average expression by cells in the cluster against the average expression by all cells not in the cluster. DEGs with an average log-fold change higher than 0.25 were used for further analysis. Gene set enrichment analysis (GSEA) showed that cluster 1 had many DEGs associated with the atypical B cell (atBC) phenotype (Portugal et al., 2015; Sullivan et al., 2015), such as *FCRL5, FCRL3, ITGB2, ITGAX, TNFRSF1B, LILRB1, CD19* and *MS4A1* (Figure 1D; Figure S1A-B). Cluster 2 expressed the lymphoid homing gene *CCR7* and the antiproliferative *BTG1* gene (Figure 1D; Figure S1B) suggesting that this may be a quiescent resting/central B cell memory subset and therefore were classified as memory B cells (MBC), while GSEA analysis revealed that these cells had a transcriptional profile similar to naïve B cells (Figure 1D; Figure S1A). Cluster 3 expressed high levels of *CXCR3* and was found by GSEA to be somewhat enriched for genes associated with previously described activated B cells (ABCs) (Ellebedy et al., 2016) including high expression *CSK* and *CD52*, however a similar level of ABC gene enrichment was also seen in the cluster 1 (Figure 1D; Figure S1A-B) therefore we did not assign a designation to these cells.

Because cells were index sorted prior to sequencing, we could also measure the surface protein expression on each cell, allowing us to investigate the expression of CD27 and CD21, the markers traditionally used to distinguish different B cell types in humans. Strikingly, only 44.7% of cells in the atBC cluster had the CD27^−^ CD21^−^ phenotype typically used to describe atBCs. Similarly, only 41.2 % of cluster 3 cells had the CD27^+^ CD21^−^ phenotype of activated B cells. Finally, 37.5 % of MBCs were CD27^+^ CD21^+^ suggesting that there may be distinct transcriptional signatures that suggest greater heterogeneity than using the canonical cell surface markers used to delineate memory B cell subsets (Figure S1C-D).

We have previously reported V(D)J sequences for the adult cells reconstructed using VDJpuzzle software (Aye et al., 2020; Rizzetto et al., 2018) we further extended this analysis to cells from the children analyzed in this study to V(D)J and isotype sequence from a total of 121/163 cells. This analysis revealed that all populations, including MBCs had undergone somatic hypermutation (SHM), though levels of SHM were slightly lower in MBCs, and – consistent with previous reports (Aye et al., 2020; Murugan et al., 2018; Tan et al., 2018) – CSP-specific B cells (Figure 1E). This analysis further revealed no strong association between antibody subclass and any population of memory cells (Figure 1F). In contrast when we subdivided the populations by antigen specificity we found that PfCSP-specific B cells were predominantly atBCs which may be consistent with continuous exposure to low levels of this antigen via repeated *P falciparum* infections (Figure 1G; Figure S1E). Surprisingly – given the association of malaria exposure with atypical B cells – most PfMSP1 specific cells mapped to cluster 3 rather than the atBC population; finally, Tetanus Toxoid specific cells were mostly MBCs which is consistent with the absence of ongoing antigenic exposure (Figure 1G; Figure S1E). Overall our data on antigen-specific cells enables us to identify 3 distinct populations of circulating B cells.

### High throughput single-cell analysis identifies atBC, MBC and ABC populations in both malaria-exposed and non-exposed donors

We next wanted to know if the 3 subsets of circulating B cells we identified were specific to malaria or reflected B cell memory in general. Moreover, we were concerned that the association of antigen with cell populations, while striking, could be a result of cells of different specificities coming from different donors and thus our analysis might be confounded by batch effects (Figure S1E-F). Finally, we wanted to sample a larger number of cells as the relatively small number of cells analyzed may not have allowed us to discern smaller populations of B cells. We therefore used the 10x Chromium platform to sequence single CD20^+^ CD19^+^ IgD^−^ memory B cells, regardless of antigen specificity, sorted from the PBMCs of two non-exposed donors (Non-Exp) and two malaria-exposed (Malaria-Exp) donors (Table S1). We also included barcoded antibodies specific for CD11c, CXCR3, CD21 and CD27 to perform Cellular Indexing of Transcriptomes and Epitopes by sequencing (CITE-seq) analysis linking surface protein expression to transcriptomic data (Stoeckius et al., 2017). We chose these markers based on our Smart-seq2 experiment and to reconcile our data with established markers. Finally, we used single cell immune profiling to obtain paired heavy and light chain V(D)J chain sequences for the BCR of each individual B cell.

After quality control steps, for the malaria-exposed donors, 1448 (Malaria-Exp 1) and 5719 (Malaria-Exp 2) cells with median genes per cell of 1576 and 1652 respectively were sequenced. While in the non-exposed individuals a further 2252 (Non-Exp 1) and 3561 (Non-Exp 2) cells were sequenced with median genes per cell of 1668 and 1535 respectively. Unsupervised clustering using *Seurat* was performed on each sample to identify any non-B cell clusters, to be removed before combining samples together (Figure S2A). Strikingly, in one of the exposed individuals (Malaria-Exp 1), we identified a cluster enriched for *CD5* and *BCL2* in which all cells expressed the same heavy and light chain immunoglobulin genes (*IGHV7-81* and *IGKV1-8)*. We concluded that these cells might be from a premalignant B cell clonal expansion and were subsequently removed from further analysis (Figure S2A).

Following removal of non-B cell populations, we used *Seurat’s* integration feature to remove batch effects between samples and combine all 4 into one integrated dataset (Figure 2A). Unsupervised clustering was then performed on this combined dataset of 12 621 cells, which revealed 11 conserved clusters (Figure 2A; Figure S2B). Three of these clusters which were more distantly related to the others appeared to correspond to naïve B cells, PCs and a population of cells expressing high levels of proliferation markers (Figure 2B; Figure S2B). The PC cluster could be discerned by the high expression of the transcription factors *XBP1, IRF4* and *PRDM1* (Figure S3A), which are all associated with controlling PC differentiation and maintenance (Klein et al., 2006; Reimold et al., 2001; Shaffer et al., 2002). The naïve cluster was characterized by expression of *IGHD*, as well as *BACH2* and *BTG1* (Figure S3A) which are transcriptional repressors associated with cellular quiescence (Guehenneux et al., 1997; Muto et al., 1998; Tsukumo et al., 2013). The third cluster we designated proliferating (Prol) cells, due to their high expression of *CD69, IRF4, MYC* and *CD83* (Figure S3A).

**Figure 2:**
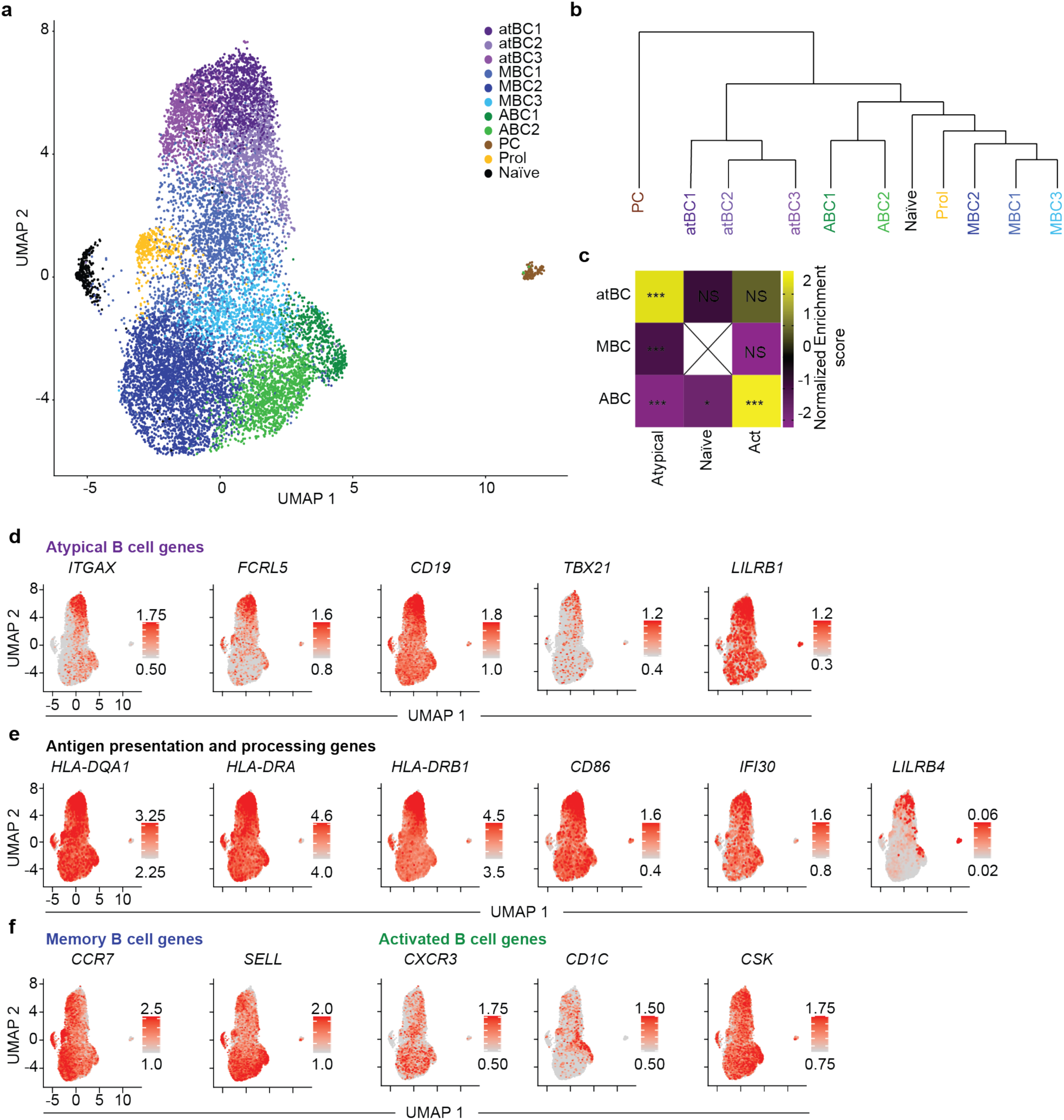
High throughput single cell analysis of reveals the full diversity of circulating B cell populations. Single B cells were sorted from 2 malaria exposed Kenyan individuals and 2 Australian individuals and gene expression was assessed using 10x chromium methodology **A**. Unsupervised clustering of circulating mature IgD^−^ B cells pooled from all individuals visualized using UMAP. Each cell is represented by a point and colored by cluster. **B**. Phylogenetic tree based on the ‘average cell’ from each cluster showing relationships in gene expression patterns between clusters. **C**. Heatmap displaying the normalized enrichment scores of multiple GSEA comparing each cluster against previously published gene sets. **D, E & F**. Expression of atBC (D), antigen presentation (E), MBC and ABC (F) genes projected onto UMAP plots. Color was scaled for each marker with highest and lowest log-normalized expression level noted. Where the exact p value is not quoted * p<0.05, **p<0.01, ***P<0.001.

DEGs were again identified in the same manner as with the Smart-seq 2 dataset. Visual inspection of a heatmap showing the top DEGs combined with phylogenetic analysis (Figure 2B; Figure S2B) suggested that the remaining 8 clusters could be grouped into 3 distinct “superclusters”. Similar to the 3 clusters identified in our Smart-seq 2 analysis, GSEA showed that the 3 superclusters identified using the 10x Chromium correspond to atBC, MBC and ABC populations (Figure 2C; Figure S3B). Notably the ABC cells identified in the 10x Chromium analysis appeared to express a stronger ABC signature than the “cluster 3” cells from the Smart-seq2 analysis. To determine the relationship between our Smart-seq 2 clusters and those found using 10x Chromium, we combined both datasets using *Seurat’s* integration command (Figure S3C-D). This integrated dataset revealed that there was general consensus between the atBC and MBC clusters identified separately using the Smart-seq 2 and 10x Chromium methodologies. However only ∼20% of the “cluster 3” Smart-seq2 cells were found to be ABCs in the integrated dataset. Rather, these cells clustered more closely with the MBC1 subpopulation within the MBC supercluster (Figure S3C-D).

Similar to the Smart-seq2 atBCs, cells in the 10x Chromium atBC “supercluster” showed higher expression of atypical genes such as *ITGAX, FCRL5, TBET, LILRB1* and *CD19* (Figure 2D; Figure S3B). The presence of the three sub-clusters (designated atBC1, atBC2 and atBC3) showed that there was nonetheless some heterogeneity within this population, such as the lower expression of *ITGAX, FCRL5* and *TBET* in atBC2 and almost no expression of these markers in atBC3. In agreement with observations seen in mouse models suggesting that these cells are primed for antigen presentation (Rubtsov et al., 2015), we found that cells from the atBC supercluster upregulate genes associated with antigen presentation and processing (Figure 2E). The MBC supercluster, made up of the sub-clusters designated MBC1, MBC2 and MBC3, lacked a clear core gene signature with less than 10 positively expressed DEGs suggesting that these cells were in a state of quiescence. Consistent with the idea that these cells represent a recirculating, memory population, many were found to have high mRNA expression of the lymphoid homing receptors *CCR7* and *SELL* (Figure 2F). The ABC super-cluster, made from 2 clusters (ABC1 and ABC2), had high expression of the activated B cell genes such as *CD1C* and *CSK* (Figure 2F). *CXCR3* was not highly expressed in the ABC supercluster, but rather was most abundant on the MBC1 population, which is consistent with the fact that most of the Cluster 3 *CXCR3* expressing cells identified in the Smart-seq2 analysis map to this population.

### Pseudotime Analysis reveals two distinct lineages of circulating B cells

To determine the lineage relationships between the distinct populations identified by unbiased hierarchical clustering we used pseudotime analysis using the R package *Monocle 3* (Trapnell et al., 2014). Visual inspection of the resulting UMAP appeared to reveal 2 distinct, major branches of circulating B cells (Figure 3A). The first branch, made up of the more “classical” MBC2 and MBC3 memory populations, and ABCs, had low progression along pseudotime (Figure 3A), indicating they were more closely related to naïve cells, which marked the beginning of pseudotime. Strikingly, pseudotime appeared to form a loop, indicative of cells transitioning between states, such as activated (ABC1 and ABC2) and quiescent (MBC2 and MBC3) rather than forming terminally differentiated populations. The second, more “atypical” branch consisting of the atBCs, MBC1 and some MBC3s, had progressed further along pseudotime, suggesting they had differentiated further away from naïve precursors than had the “classical” branch. Pseudotime could again be seen to form a loop, suggesting that atBCs may also fluctuate between resting (MBC1 and MBC3) and activated (the atBCs) states. The proliferating cells appeared to form their own distinct branch, or lineage, separate from either ABCs or atBC. Interestingly, The PCs appeared entirely detached from the pseudotime pathway, indicating that an intermediate PC population could not be found amongst circulatory B cells.

**Figure 3:**
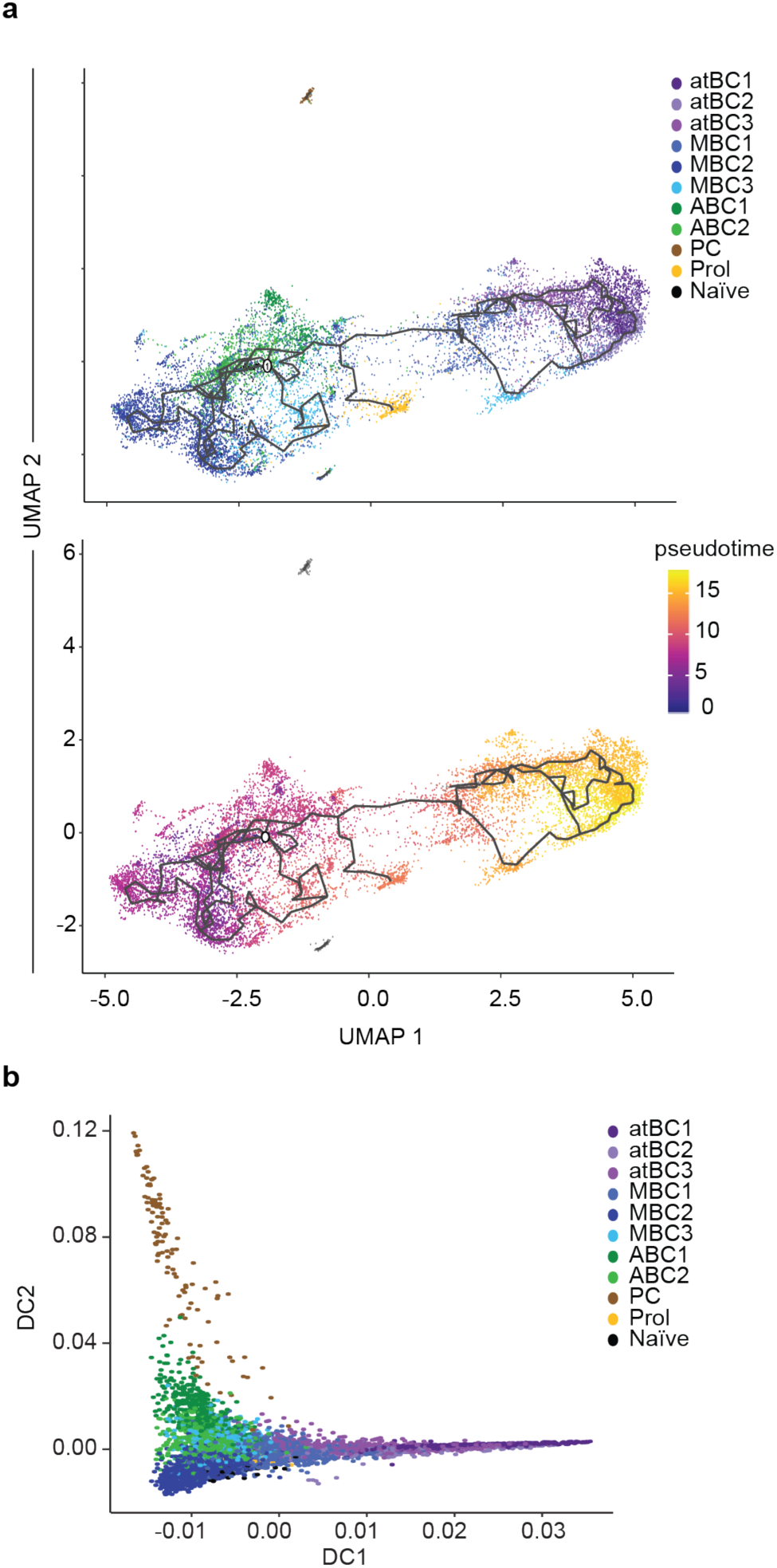
Pseudotime analysis and diffusion mapping reveal 2 distinct lineages of circulating B cells. **A**. Pseudotime analysis of circulating B cells generated visualized using UMAP. Each point represents a cell and is colored by cluster or progression along pseudotime. **B**. Diffusion map showing diffusion components (DC) 1 and 2, each cell is represented by a point and colored by cluster.

Similar results were obtained when we used a diffusion mapping approach to examine lineage relationships. Two distinct branches of cells could be seen diverging from a central cluster of MBCs (Figure 3B). atBCs formed the first branch with atBC2s and atBC3s predominantly found at the base of the branch, closer to the MBCs and atBC1s found at the tip. The second branch was made of ABCs at the base extending to PCs at the tip suggesting that following immunization or infection, MBCs may follow either a classical activation pathway that ultimately leads to terminally differentiated antibody secreting PCs or an “atypical” pathway, culminating in the formation of Tbet^+^ FCRL5^+^ CD11c^+^ atBCs.

### Atypical B cells are represented at high frequencies in all individuals, but are not necessarily CD27^−^ CD21^−^

Having identified 11 populations of circulating B cells by transcriptional profiling, we verified that all populations could be found in all individuals tested, albeit in slightly different proportions (Figure 4A; Figure S4). The malaria-exposed donors had high numbers of atBC1 and atBC2 populations, but surprisingly the non-exposed donors also had significant numbers of atBCs (∼20%) that were largely atBC3 (Figure 4A; Figure S4B). Non-exposed donors and malaria-exposed donors alike also additionally carried high numbers of the MBC1 population (∼20%) which appears related to the atBC lineage (Figure 4A; Figure S4). This number of atBCs and related cells, as identified by transcriptomic techniques, contrasts with previous flow cytometry analysis which shows that non-malaria exposed healthy donors typically carry few (generally <5%) CD27^−^, CD21^−^ atypical cells (Illingworth et al., 2013; Weiss et al., 2009).

**Figure 4:**
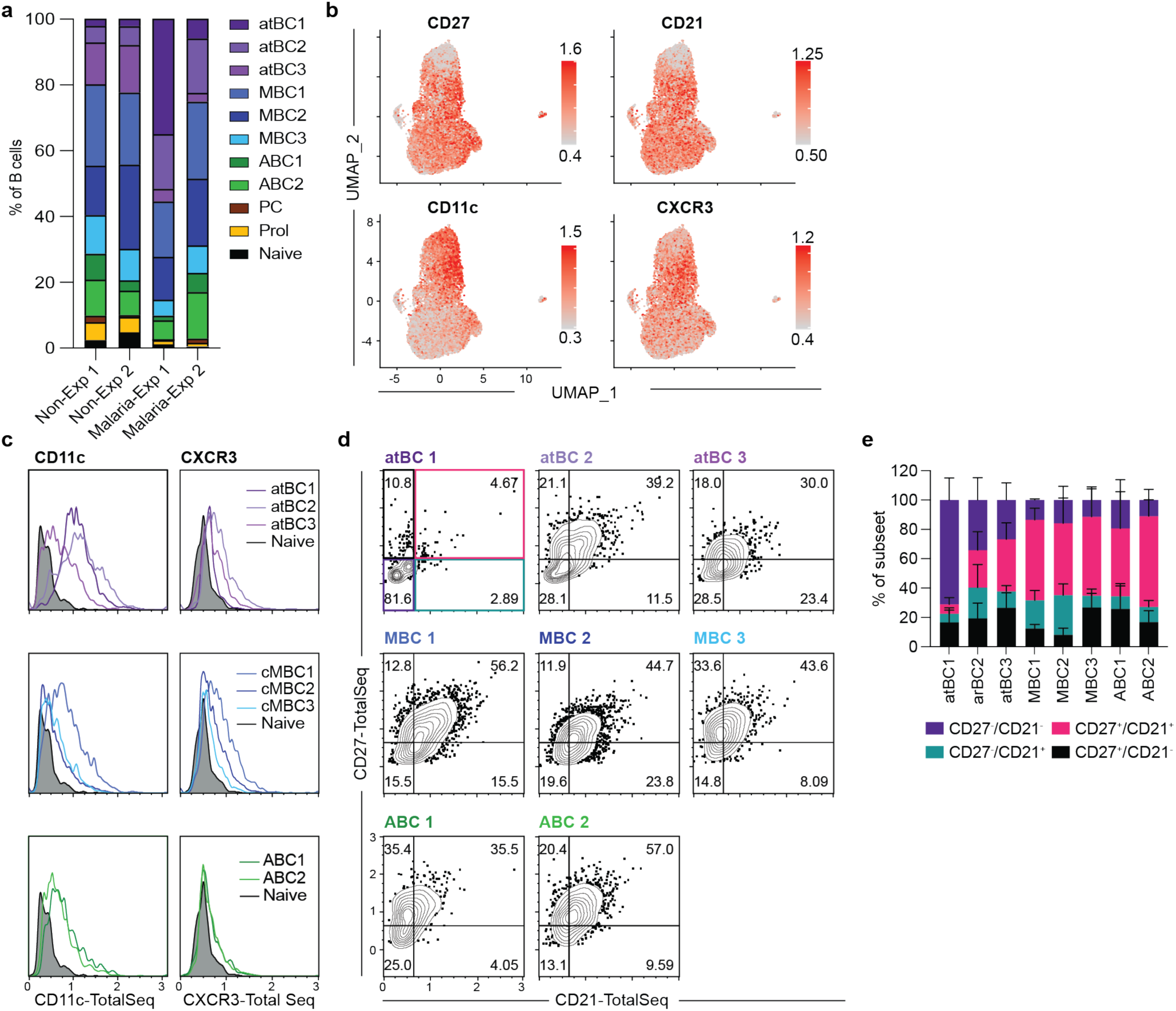
CITE-Seq analysis reveals a cryptic population of atBCs found predominantly in Australian individuals. CITE-seq analysis to correlate expression of cell surface markers with gene-expression was performed on cells from the four donors described in figure 2 **A**. Percentage of cells from each individual found in each cluster. **B**. Surface protein expression measured by CITE-seq projected onto UMAP plots. Color was scaled for each marker with highest and lowest log-normalized expression level noted. **C**. Histogram plots showing the expression of CD11c and CXCR3 for the different atBC, MBC and ABC clusters, grey histogram represents expression on naïve B cells; data are concatenated from all individuals. **D**. Contour plots showing the expression of CD27 and CD21 as measured by CITE-seq; data are concatenated from all individuals E. Quantification of (D), data show the mean proportion per individual ± s.d..

To address this discrepancy, we used CITE-seq to correlate the cell surface levels of our candidate markers CD11c and CXCR3 as well as CD27 and CD21 to our transcriptomic data for each cell and cluster (Figure 4B). CITE-seq data was exported into flow cytometry analysis software (FlowJo) for further processing and presentation. CD11c was abundant on atBC1 and atBC2 cells which are common in malaria exposed individuals but was only found at low levels on the atBC3 population which was preferentially found in the non-exposed donors (Figure 4C). Among MBC populations we found that CD11c was more abundant on the MBC1 populations that appears related to atBC lineage. Analysis of CD21 and CD27 expression showed that while atBC1 cells were almost exclusively CD21^−^ and CD27^−^, the other populations of atBCs had more heterogeneous expression of these markers (Figure 4D and E). Thus, the atBC3 population appears to be a cryptic atBC population which cannot be detected via conventional flow cytometry strategies. These data may support the conclusion from our pseudotime analysis that there is a spectrum of activation within atBCs, with CD21^−^, CD27^−^, FCRL5^+^, CD11c^+^ B cells representing a more activated phenotype (atBC1 and 2) while other cells become more quiescent, losing expression of these markers while retaining a core gene signature (atBC3 and MBC1).

We further used this analysis to investigate the utility of our markers for identifying MBC and ABC populations. ∼50% of the MBC1 and MBC2 populations were CD27^+^, CD21^+^, but only ∼20% of the ABC populations resembled the activated CD27^+^, CD21^−^ phenotype. MBC3 also looked somewhat activated as ∼30% of cells were CD27^+^ CD21^−^ (Figure 4D and E). This is consistent with the observation that this population does appear to express some activation genes, albeit at low levels (Figure 2F-Figure S2). As expected from our gene expression data CXCR3 was not a useful marker for the ABC populations, rather CXCR3 was most abundant on MBC1 cells consistent with the *CXCR3*-expressing “cluster 3” cells from the Smart-seq2 data set mapping to this population. CXCR3 was also found on the atBC2 population which is consistent with this being a marker of “activated” atBCs. Overall these data show that that CD27 and CD21 poorly mark different B cell memory subsets, however they suggest that – while imperfect – CD11c is a useful maker of the atBC lineage.

### Subsets of circulating B cells do not segregate with Ig subclass or V region usage

In murine models different memory B cell subsets may be defined by their Ig-subclass, including in malaria infection (Krishnamurty et al., 2016; Pape et al., 2011). We therefore performed analysis of the VDJ and constant region usage of the heavy chains of our different B cells populations. The switched subclasses (IgM, IgG1, IgG2, IgG3, IgG4, IgA1, IgA2 and IgE) were found in all non-naïve B cell populations, with the exception of IgE, of which only 2 cells were found across all donors (Figure 5A). The population we identified as naïve was largely IgM^+^ with some cells expressing IgD^+^ further supporting this designation. In both malaria-exposed donors, IgG3 was overrepresented in the atypical memory B cell compartment, which is consistent with previous reports (Knox et al., 2017; Obeng-Adjei et al., 2017). Overall, these data suggest that the transcriptional signatures are distributed across all Ig subclasses.

**Figure 5:**
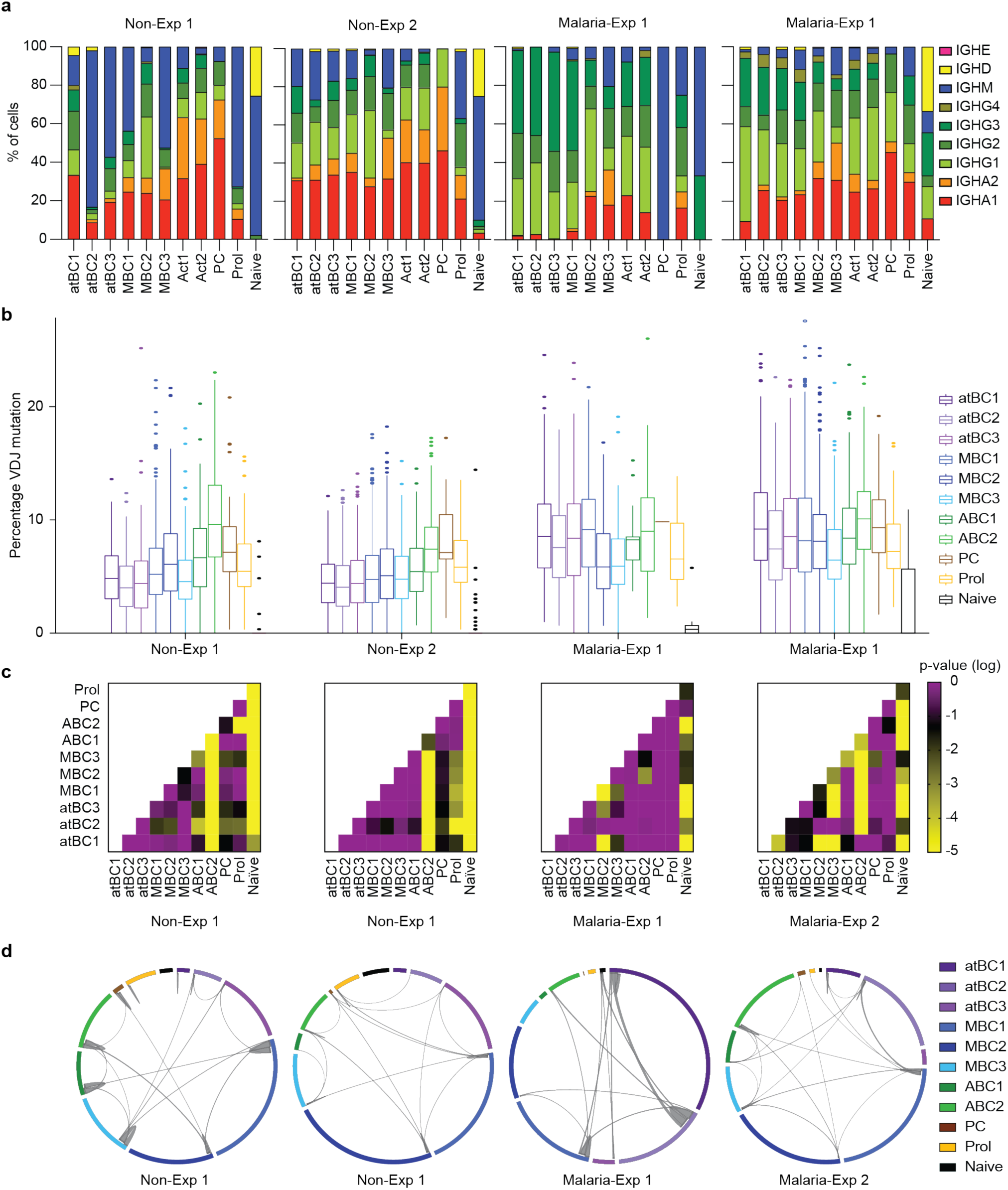
Lack of association between BCR variable and constant regions with different B cell subsets. V(D)J and constant region sequences for each cell from each donor described in figure 2 were mapped to the individual transcriptomes and relationships analyzed **A**. percentage of isotype usage for each cluster per individual. **B**. Percentage of mutations found in the heavy chain V(D)J region of cells for each cluster in each donor, mean ± s.d. shown. **C**. Heatmaps displaying the pairwise p-values from Tukey’s post-tests based on one-way ANOVA of the data in (B) to determine the association between cell type and mutation frequency with subclass and individual also included in the model as fixed factors. **D**. Circos plots showing clonal B cell populations per individual, the thickness of the lines between or within clusters denotes the number of cells that belong to shared/expanded clones.

We further examined the variable region sequence of each cell to determine the level of SHM and clonal relationships based on identical *IGHV* CDR3’s with matching *IGLV or IGKV*. Similar to the constant region, V region usage was similar between all donors and B cell clusters (Figure S5A-B). All clusters, with the exception of the “naïve” cells showed significant SHM, confirming that these cells were antigen experienced, post-GC B cells (Figure 5B). The degree of SHM differed significantly based on the cell population and donor (Figure 5B). Notably across all populations, non-exposed donors apparently carried lower levels of SHM than malaria-exposed donors, perhaps indicating lower lifetime pathogen burden (Figure 5B). However, most V(D)J databases are based on Europeans and may reflect allelic differences between populations of African and European ancestry. In 3/4 donors, ABC2 and PCs had higher levels of SHM compared to either atBC populations or MBC populations (Figure 5B-C), which may be consistent with these populations being related as indicated by diffusion mapping analysis (Figure 3C). Finally, 1-5% of all BCRs sequenced were shared between 2 or more cells in each sample (Figure 5D). In all individuals, expanded clones could be found, in most cases these expanded clones were found within clusters, however clones could also be found shared between clusters, including across superclusters indicating that a single clone can potentially adopt multiple cell fates (Figure 5D).

### Flow cytometry analysis reveals heterogeneity in atBC populations from malaria exposed individuals

To extend the analysis of circulating B cells beyond the original 4 donors, we performed flow cytometry analysis of B cells from the PBMCs of 11 malaria-exposed and 7 non-exposed individuals (Table S1). Because the pseudotime analysis suggested that both classical and atypical lineages can cycle between activated and resting states we also included CD71 as a marker of B cell activation (Ellebedy et al., 2016). To identify atBCs we used a panel of CD11c, FCRL5, CD27, CD21 and T-bet, though none of these markers would be likely to capture the “cryptic” atBC3 population found in non-exposed healthy adults which did not express any obvious candidate surface markers based on our transcriptomic and CITE-Seq analysis.

Consistent with our own transcriptomic and CITE-seq data as well as the data of others (Portugal et al., 2015; Weiss et al., 2009), we found that CD11c^+^ cells were considerably enriched in malaria-exposed donors (Figure 6A-B). We were able to identify a small population (∼5%) of CD11c^+^ B cells among the non-exposed donors which were CD19^hi^, CD20^hi^ and FCRL5^hi^, but only a few of these cells were CD21^−^, CD27^−^ T-bet^hi^ (Figure 6C-E). The CD11c^+^ cells from malaria-exposed donors expressed high levels of T-bet and were mostly CD21^−^ CD27^−^ (Figure 6D-E). This supports the finding that there may be a spectrum of atBCs, which may also explain why these cells have been identified and characterized in slightly different ways in different pathologies (Jenks et al., 2018; Moir et al., 2008; Weiss et al., 2009). In both non-exposed and malaria-exposed donors around 30% of B cells were CD71^+^ CD11c^−^ (Figure 6A-B) which is consistent with the proportion of ABCs identified by single cell RNA-seq. However, CD71 was also expressed on a high proportion of the CD11c^+^ B cells. This further supports the observations from the pseudotime analysis that atypical cells may fluctuate between activated and resting states. Further analysis found that these CD71^+^ atBCs expressed high levels of CXCR3 (Figure 6E), further supporting the observation from CITE-seq data that CXCR3 is a marker of activated atBCs.

**Figure 6:**
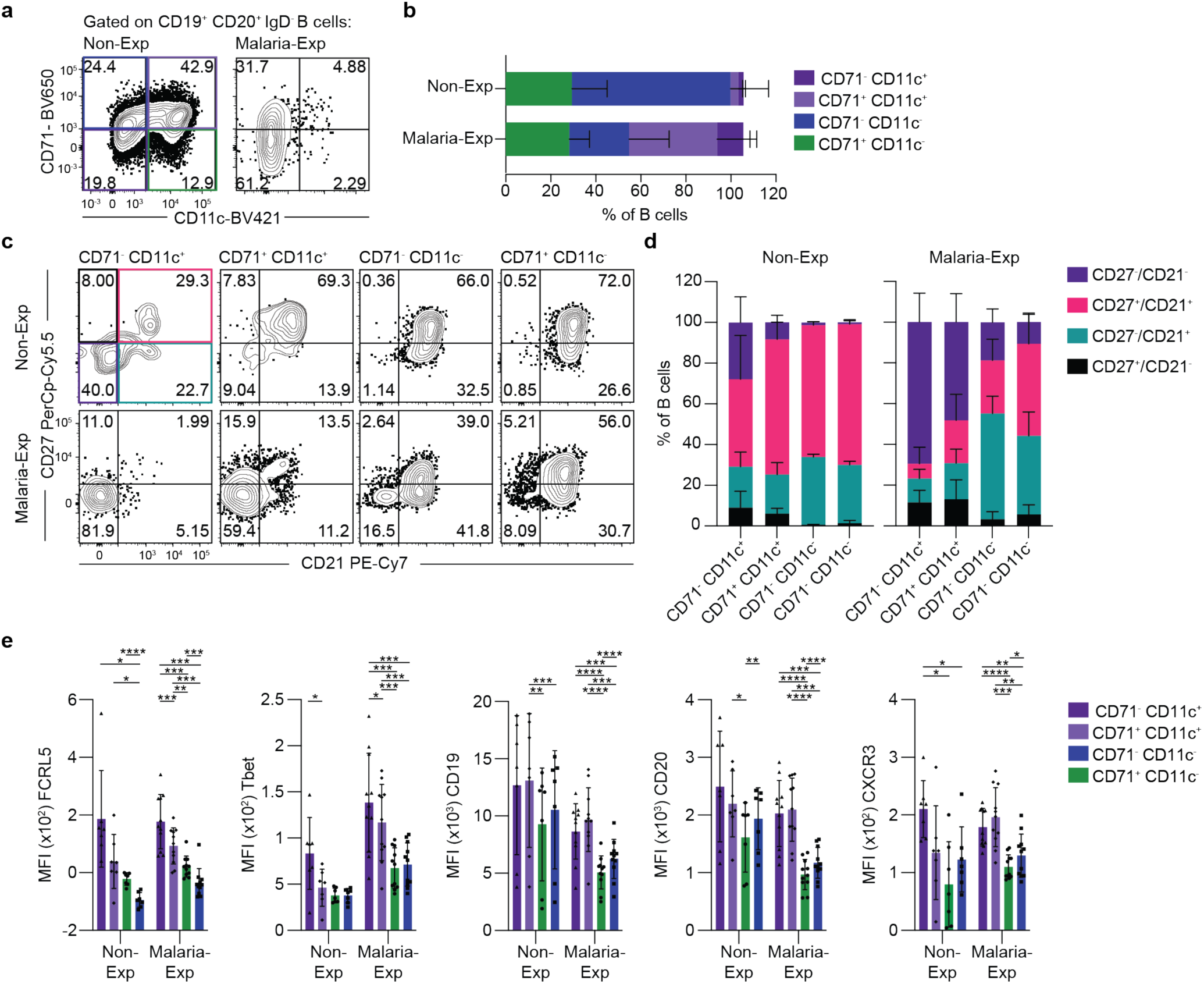
CD11c and CD71 identify atBC, ABCs and MBCs via flow cytometry. PBMCs from 7 Non-Exp and 11 Malaria-Exp donors were isolated and analyzed by flow cytometry for expression of markers associated with different B cell populations **A**. Flow cytometry plots from representative individuals showing the CD11c and CD71 expression on mature IgD^−^ B cells. **B**. Quantification of (A) showing the percentage of cells found in each cell type by country, bars represent mean ±s.d.. **C**. Representative flow cytometry plots showing the expression of CD27 and CD21 per cell type. **D**. Quantification of (C) showing the percentage of cells separated by expression of CD27 and CD21 found in each cell type, bars represent the mean proportion ±s.d.. **E**. The expression of surfaces markers on each cell type, measured by MFI, analysis was done using 2-way ANOVA, bars represent mean ±s.d..

### CD11c^+^B cells arise in the primary response and respond to booster immunization

Our pseudotime analysis indicated that the majority of circulating B cells can be separated into two distinct lineages, classical or atypical, with cells in each lineage able to fluctuate between activated and resting states. While the diffusion map results suggest that the classical lineage gives rise to antibody secreting PCs, the role of the atypical lineage in the immune response is still not understood. In the past it has been suggested that atBC are an exhausted or dysfunctional cell population that arise following chronic antigen exposure (Portugal et al., 2015; Sullivan et al., 2015). We therefore wanted to examine a situation in which we could (i) track antigen specific cells after primary exposure and (ii) continue to follow those cells upon antigen re-exposure. We would thus be able to determine the point in an immune response at which atBCs arise as well measure their longevity and their capacity to be recalled. To meet these criteria, we tracked B cells specific for PfCSP in a cohort of malaria-naïve individuals who were given three doses of a whole *P. falciparum* sporozoite vaccine (PfSPZ) at 8-week intervals (Ishizuka et al., 2016; Lyke et al., 2017). These sporozoites are irradiated so do not establish ongoing infection, though there is previous evidence of antigen persistence (Cockburn et al., 2010). Blood samples were obtained at the time of each vaccination and 1 wk and 2 wks post each vaccination and flow cytometric analysis was performed to identify and characterize PfCSP-specific B cells (Figure 7A; Figure S6A). We were further able to identify PfCSP specific PCs in these individuals (Figure S6A).

**Figure 7:**
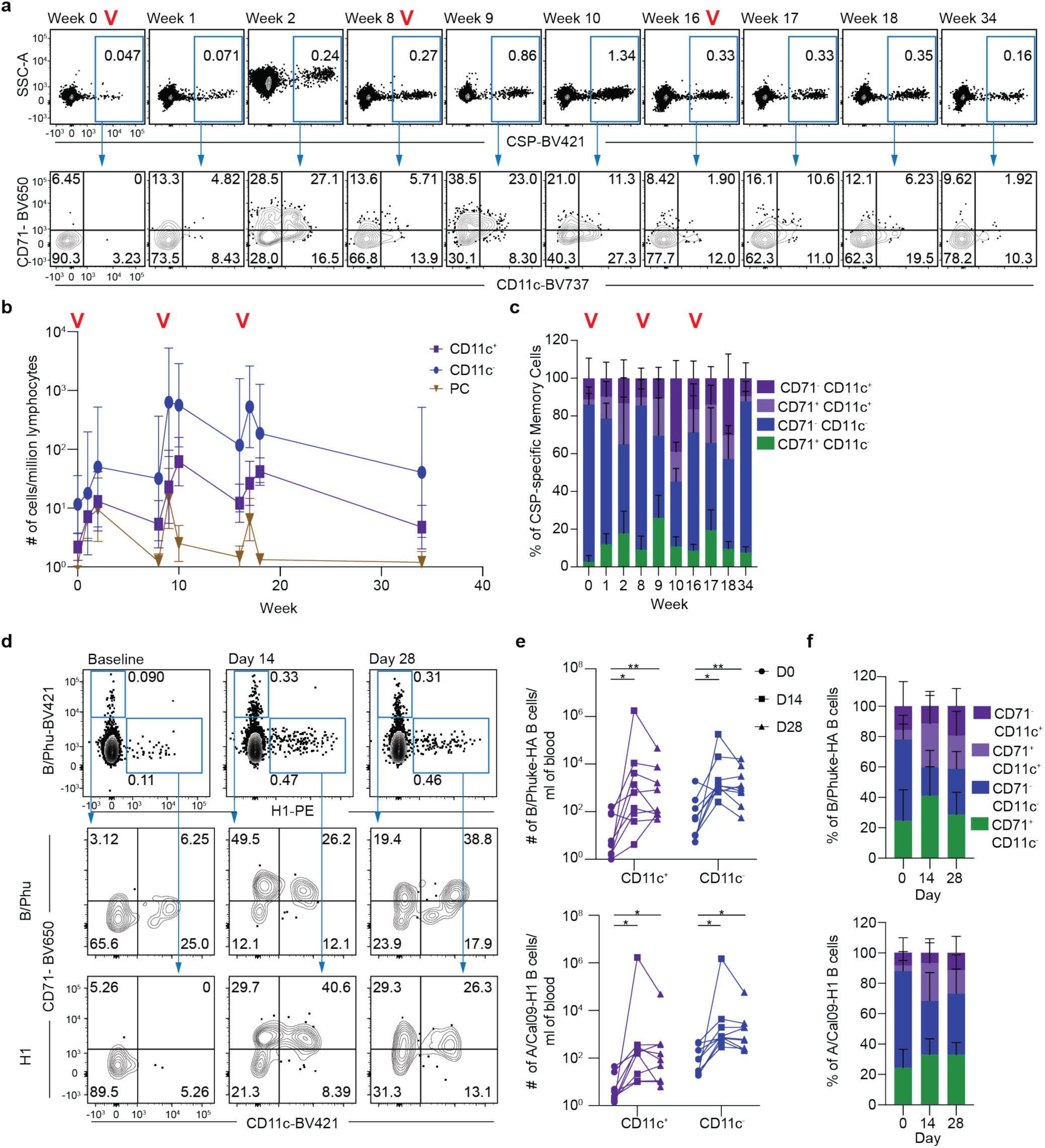
Antigen-specific atBCs arise during the primary response and can be effectively recalled. **A**. 15 individuals were vaccinated with 3 doses of 9 × 10^5^ PfSPZ at 8-week intervals, with blood drawn at the indicated timepoints; panel shows representative flow cytometry plots from a single individual of the gating of CSP-specific IgD^−^ B cells and the CD71 and CD11c expression found in these cells over time. Red “V”s indicate time points where booster immunizations were given. **B**. Kinetics of the CSP-specific B cell response quantified by the number of cells per million lymphocytes. **C**. The percentage of CSP-specific memory cells divided by CD11c and CD71 expression over time **D**. 9 individuals were vaccinated with inactivated influenza vaccine (IIV) with blood drawn at baseline and 14 or 28 days later; panel shows representative flow cytometry plots showing the number of either B/Phuket or H1-specific IgD^−^ B cells and the CD71 and CD11c expression found in these cells over time. **E**. Kinetics of the influenza-specific B cell response quantified by the number of cells per mL of blood. **F**. The percentage of influenza-specific memory cells divided by CD11c and CD71 expression.

By 1 week after immunization the number of CSP specific B cells had begun to increase, though most remained CD71^−^, CD11c^−^ (Figure 7A-C). However, by 2 weeks post immunization, significant populations of CD71^+^ and CD11^+^ B cells could be seen, with many of the CD11c^+^ B cells co-expressing CD71, matching those seen in malaria-exposed individuals (Figure 7A-C). By 8 weeks post immunization distinct CD71^+^ and CD11c^+^ B cell populations were seen, but the CD11c^+^ B cells no longer expressed high levels of CD71 further supporting the idea that this is a marker of recent activation on atBCs (Figure 7C). Importantly, both CD71^+^ and CD71^−^ CD11c^+^ B cells expressed high levels of CD19, CD20 and low levels of CD21 and CD27 further supporting the fact that these cells represent a bona-fide atBC population (Figure S6B-C).

We further tracked these B cell populations following boosts after 8 and 16 weeks (Figure 7A-C). There was a significant expansion of the all B cell populations at the first boost, while magnitude of the response did not change at the second boost. During the first boost the CD71^+^ CD11c^−^ population peaked after 1 week; more rapidly than during the primary response (Figure 7C), however the total CD11c^+^ population continued to expand and only peaked 2 weeks after each boost. Notably the kinetics of the atBC population are distinct from those of the PC population which is only detectable 1 week after each boost, which further suggests that in normal conditions atBCs are not pre-PCs. Finally, 16 weeks after the final boost (wk 34) we examined the number and phenotype of the different cell populations. At this “memory” timepoint most cells were double negative, but residual populations of CD11c^+^ and CD71^+^ B cells could still be detected (Figure 7A-C).

Finally, we wanted to know if our findings could extend beyond *Plasmodium* to another vaccination setting, also involving acute exposure to antigen. We therefore examined how these cells responded in a recall response to the inactivated influenza vaccine (IIV) (Koutsakos et al., 2018). Using recombinant rHA probes to two IIV antigens: A/California/07/09^−^H1N1 (A/Cal09^−^H1) or B/Phuket/3073/2013 (BHA; Yamagata lineage) we were able to identify influenza specific B cells from the peripheral blood of subjects prior to immunization (Figure 7D; Figure S6D). CD71^+^ ABCs, CD11c^+^ atBCs and double negative MBCs to both influenza antigens could be found at baseline in most individuals as expected, reflecting past exposure to influenza virus infection or vaccination (Figure 7D-F). Following immunization, CD71^+^ and CD11c^+^ cells expanded alongside the double negative population. CD71^+^ CD11c^+^ and CD11c^−^ B cells peaked first, in this case 2 weeks after immunization particularly against the B/Phuket-HA strain, while CD11c^+^ B cells had a more sustained expansion (Figure 7D-F). Again, influenza specific CD11c^+^ cells found after vaccination expressed high levels of CD19, CD20 and FCRL5, and high proportions of these cells were CD21^−^ and CD27^−^ consistent with them being an atypical population (Figure S6E-F). All together these data reveal that atBC arise during the primary response and that these cells appear to participate in recall responses upon re-exposure to antigen, countering the suggestion that these cells are a dysfunctional or exhausted population.

## Discussion

Atypical B cells, which have been conventionally defined based on specific cell surface markers have been found in excess in many pathological conditions, in particular chronic or repeating infections and autoimmunity. This association with disease, along with the difficulty of re-stimulating these cells *in vitro* has led to the assumption that these are an exhausted or even pathological population. Our observations using single cell transcriptional analysis that atBC are more abundant than expected in both healthy non-exposed and malaria-exposed individuals lead us to test the hypothesis that these cells are a stable lineage that arise in response to antigenic stimulation. Accordingly, we tracked these cells in controlled conditions of vaccination and further showed that these cells arise during the primary immune response and can be recalled normally on multiple re-exposures. Importantly these cells are not merely recently activated B cells as they have a distinct gene signature from previously described ABCs. Thus, while – in accordance with previous literature – we have used the term “atypical” to describe this lineage, the data presented here suggests that atBCs contribute to antigen-specific primary and recall antigen specific responses, and thus are a typical part of the B cell response to antigen.

It has been hypothesized that atBCs are exhausted (Portugal et al., 2015; Sullivan et al., 2015) or recently activated cells (Jenks et al., 2018; Perez-Mazliah et al., 2018). However, our observations that atBC are generated during the primary response to sporozoite vaccination and can be recalled following booster immunizations, either to sporozoite or influenza antigens would suggest that atBC are not necessarily exhausted cells and can in fact participate in normal immune responses. Furthermore, our single cell RNA-seq data clearly differentiates atBC from previously described ABCs. Nonetheless there is also evidence that atBCs themselves can be activated which may resolve some of the discrepancies seen in the literature. Specifically, we have shown that CD71^+^ atBC appear early in the response and abate quickly, giving rise to a more conventional CD71^−^ atBC population. These CD71^+^ atBCs likely represent a population of recently recalled cells. Interestingly, while the first study to describe ABCs cells used CD71 as the primary marker for ABC (Ellebedy et al., 2016), CD11c^+^ B cells, which we have shown can express CD71 were not excluded. Thus, this bulk sorted population may have contained some atBCs which may explain why in our smaller Smart-seq2 dataset, the atBC cluster was found to be enriched for ABC genes. We did however observe that the antigen-specific atBC population does diminish overtime in our sporozoite vaccinated individuals. One explanation for this is that atBCs are not a bona fide memory population that are as long-lived as the MBCs. However, the presence of a sizeable CD11c^−^ FCLR5^−^ CD27^+^ atBC (atBC3) as well as MBC1 populations in healthy non-exposed individuals suggests that there may exist a pool of “cryptic”, quiescent MBCs primed to differentiate into CD11c^+^ atBCs following re-exposure.

We initially described a tripartite division in our description of the circulating B cell subsets. This is based on the unsupervised clustering of our single cell RNA-seq datasets revealing distinct “superclusters” of circulating B cells: Activated (ABC), Memory (MBC) and Atypical (atBC). However, an alternative classification informed by pseudotime analysis and our data from vaccination cohorts would suggest a division of two distinct lineages, or pathways of MBCs and their corresponding activated populations. For example, in the classical pathway, MBC2 are quiescent cells that, following antigenic stimulation, transition into proliferating ABCs that can go on to either terminally differentiate into PCs or reseed the MBC pool. A previous study showed that up to 60% of ABC clones sequenced at day 7 could be found in the MBC subset 90 days post vaccination, revealing that ABCs can downregulate their activation markers and become MBCs over time (Ellebedy et al., 2016). Under this model the ABC2 and MBC3 populations likely represent a spectrum of cells that were either recently activated or entering a state of quiescence, rather than defined B cell fates. Interestingly, our VDJ analysis revealed that B cell clones could be found in both cell lineages, indicating that the progeny of a single B cell may adopt both cell fates

While our data show that atBC are part of a normal B cell response, the function of these cells remains elusive. Early studies revealed that it was difficult to differentiate atBCs into antibody secreting cells under standard conditions *in vitro* (Moir et al., 2008; Portugal et al., 2015; Sullivan et al., 2015). However bulk RNA-seq analysis of atBC-like cells from SLE patients showed higher expression of genes associated with PC maintenance in this populations lead to the hypothesis that atBCs represent a precursor PC population (Jenks et al., 2018). Our analysis however, using single cell RNA sequencing techniques, could not find any evidence of these genes being upregulated in any of our atBC populations. We also observe that our atBC population in sporozoite vaccinated individuals continues to expand even 2 weeks after booster immunization while PC populations peaked 1-week post-immunization. A reconciliation of these conflicting results may be that in pathogenic conditions such as SLE, atBCs can be driven to become pathogenic antibody secreting cells. Consistent with this the TLR7 pathway is implicated in SLE development and it has been found that including TLR7 agonists within the stimulating condition can help differentiate atBCs into PCs (Jenks et al., 2018; Perez-Mazliah et al., 2018; Rivera-Correa et al., 2019; Rubtsova et al., 2013).

A potential role for atBC may be specifically in the clearance of viral infection, as research in murine models has found that T-bet^+^ CD11c^+^ B cells, are required for effective clearance of viral infection (Barnett et al., 2016; Rubtsova et al., 2013). Our study has focused on relatively inflammatory B cell stimuli such as malaria infection and attenuated pathogen vaccines. It may be that atBC are preferentially formed in these conditions rather than subunit vaccination in less inflammatory adjuvants such as alum and thus would be expected to play a role in control of infection. Finally, it has been proposed that atBCs are potent antigen presenting cells (Rubtsov et al., 2015). In agreement with this we did find that human atBC do appear to have higher expression of MHC II as well as the co-stimulatory molecule CD86 and components of the MHC Class II antigen processing pathway, indicating that they present more antigen then other B cell types. B cells have recently been shown to be required for the priming of Tfh cells in the context of malaria infection, though the role of individuals subsets was not studied (Arroyo and Pepper, 2020). Intriguingly another mouse study also revealed that ablation of CD11c^+^ B cells lead to the collapse of germinal centers after one immunization (Baumjohann et al., 2013). Thus, these data suggest that atBCs could represent a population of specialized antigen presenters, although further investigation is needed to confirm this hypothesis.

Here we provide an atlas of the B cell subsets circulating in human blood. Our powerful single-cell RNA-seq analysis combined with CITE-seq technologies and VDJ profiling enables us to reconcile our data with existing classifications of B cell memory based on flow cytometry markers or Ig-subclass. We have been able to resolve some key controversies in the human B cell literature, most notably by finding that atBCs are more abundant than previously expected in healthy donors and showing that these cells can be induced by primary exposure to acute antigen. Thus, our data suggests that atBC are a critical and typical component of the humoral immune response.

## Author Contributions

Conceptualization (HJC, RA, AHI, FMN, IAC). Data curation (HJC, XL, TDA, FMN, IAC). Formal Analysis (HJC, RA, XL, XG, TDA, FL). Funding acquisition (KK, RAS, FMN, IAC). Project Administration (KEL, SLH) Investigation (HJS, RA, AHI, RV, EN, OK, JM, MK, THON, MN, PM, AAB, NK, SC). Methodology (AE, FL). Resources (AKW, SJK, FL, KK, RAS, FMN). Software (FL). Supervision (SJK, FL, KK, RAS, FMN, IAC). Visualization (HJS, XL, IAC). Writing – original draft (HJS, IAC). Writing – review & editing (FL, KK, RAS, FMN).

## Acknowledgements

This work was supported by start-up funds from the Australian National University to I.A.C. and NHMRC project grant support to I.A.C. (GNT1158404). We would like to thank Harpreet Vohra and Michael Devoy of the Imaging and Cytometry Facility at the Australian National University for assistance with flow cytometry and sorting. We also thank the staff of the biomolecular resource facility at the John Curtin School of Medical Research for assistance with single cell RNA-seq. Production and characterization of PfSPZ Vaccine were supported in part by National Institute of Allergy and Infectious Diseases Small Business Innovation Research Grants 5R44AI055229-11 (to S.L.H.), 5R44AI058499-08 (to S.L.H.), and 5R44AI058375-08 (to S.L.H.). We would like to thank the University of Maryland study volunteers from malaria clinical trial VRC314. We are grateful to the KEMRI/CGMRC field team for their dedication in the recruitments, malaria surveillance data and sample collection, and the laboratory team that processed the samples.

We are also indebted to the study participants. FMN was supported by an MRC/DFID African Research Leadership Award (MR/P020321/1), a Senior Fellowship from EDCTP (TMA2016SF-1513) and the samples were collected within the Kilifi immunology cohorts supported by various Wellcome grants. RA was supported through the DELTAS Africa Initiative [DEL-15-003]. The DELTAS Africa Initiative is an independent funding scheme of the African Academy of Sciences (AAS)’s Alliance for Accelerating Excellence in Science in Africa (AESA) and supported by the New Partnership for Africa’s Development Planning and Coordinating Agency (NEPAD Agency) with funding from the Wellcome Trust [107769/Z/10/Z] and the UK government. This manuscript is published with permission from the Director, KEMRI.

## Materials and Methods

### Human samples and ethics statement

All research was conducted according to the principles of the Declaration of Helsinki, which included the administration of informed consenting in the participant’s local language. Studies in Australia were further performed in accordance with the Australian National Health and Medical Research Council (NHMRC) Code of Practice.

The malaria-immunology cohort and vaccination studies, under which the samples described were collected in Kenya, were approved by the Kenyan Medical Research Institute Scientific and Ethics Review Unit, Nairobi, and the use of these samples at the Australian National University was further approved by the Australian National University Human Research Ethics Committee (protocol number 2014/102). The Kenyan adults are members of the KEMRI/Wellcome Research Programme’s longitudinal malaria immunology cohort studies in Junju and Ngerenya villages (supplementary table 1), 20 km apart from each other in Kilifi, Kenya.. In addition, we included samples from the RTS,S/AS01 phase 3 clinical trial. Blood was also drawn from healthy control Australian donors who were recruited at the Australian National University.

VRC 314 clinical trial (https://clinicaltrials.gov/; <u>NCT02015091</u>) was an open-label evaluation of the safety, tolerability, immunogenicity and protective efficacy of PfSPZ Vaccine. Subjects in the high dose cohort received a total of three doses of 9 ×10^5^ PfSPZ intravenously at week 0, 8 and 16. Blood was drawn at the time of each immunization, as well as 7d and 14 d after each immunization. Plasma and PBMCs were isolated from all samples at these timepoints. Full details of the study are described in (Lyke et al., 2017).

The investigation of B cell responses after IIV immunization was approved by the University of Melbourne Human Ethics Committee (ID 1443389.3) and the Australian Red Cross Blood Service (ARCBS) Ethics Committee (ID 2015#8). PBMCs were used from 8 donors taken on the day of IIV immunization, as well as 14 and 28 days later. Full details of the study are described previously (Koutsakos et al., 2018).

### Flow cytometry

For samples from Kenya and the IIV cohort PBMCs were thawed and washed in PBS with 2% heat-inactivated FBS. Cells were then stained with Live/Dead dye for 5 min in PBS before incubation with fluorescently labelled antibodies for a further 30 min. Details of all antibodies used are given in Table S2. Flow-cytometric data was collected on a BD Fortessa or X20 flow cytometer (Becton Dickinson) and analyzed using the software FlowJo (FlowJo). A BD FACs Aria I or II (Becton Dickinson) was used for sorting cells.

For VRC314 clinical trial specimens PBMCs were thawed into prewarmed RPMI media then washed with PBS. Cells were stained with Live/Dead dye for 15 minutes, washed in PBS with 2% heat inactivated FBS, and labelled with antibodies for an additional 30 minutes. Labelled cells were washed with PBS 2% FBS and fixed for 15 minutes in 0.5% PFA before and final wash and resuspension in PBS 2% FBS. Flow-cytometric data was collected on a BD X50 flow cytometer (Becton Dickinson) and analyzed using the software FlowJo (FlowJo).

### Tetramer Preparation

*Pf* MSP1, AMA1 and TT were biotinylated with the Sulfo-NHS-LC-Biotinylation Kit (ThermoFisher) at a ratio of 1:1 according to the manufacturer’s instructions, biotinylated (NANP)9 repeat region of *Pf* CSP was sourced from Biomatik (Ontario, Canada). Biotinylated antigens were incubated with premium^−^grade SA^−^PE and SA^−^APC (Molecular Probes) or SA-BV421 and SA-BB660 (Biolegend and BD Horizon) at a molar ratio of 4:1, added four times with 15 min incubation at room temperature.

### Single cell RNA^−^seq using Smart-seq2

Antigen-specific single cell RNA sequencing was performed using a Smart-seq 2 protocol (Picelli et al., 2014) with the following modifications. Cells were sorted into plates with wells containing 1 µl of the cell lysis buffer, 0.5 µl dNTP mix (10 mM) and 0.5 µl of the oligo-dT primer at 5 µM. We then reduced the amount reagent used in the following reverse^−^transcription and PCR amplification step by half. The concentration of the ISPCR primer was also further reduced to 50 nM. Due to the low transcriptional activity of memory B cells, we increased the number of PCR cycles to 28. cDNA was then purified with AMPure XP beads at a bead to sample ratio of 0.8:1. Sequencing libraries were prepared using the Nextera XT Library Preparation Kit with the protocol modified by reducing the original volumes of all reagents in the kit by 1/5^th^. Another round of bead cDNA bead purification was preformed using a bead to sample ratio of 0.6:1. Sequencing was performed on the Illumina NextSeq sequencing platform. Following sequencing, fastq files were passed through the program VDJPuzzle (Rizzetto et al., 2018) where reads were trimmed using Trimmomatic, then aligned to the human reference genome GRCh37 using tophat2. Gene expression profiles were then generated using cufflinks 2. As a further QC step, cells where reads were mapped to less than 30% of the reference genome were removed. All following downstream analysis for transcriptomic data was performed using *Seurat*. Details of all key reagents for single cell RNA-seq are given in Table S3.

### Single cell RNA-seq and CITE-seq using 10x Chromium

Post-sorting, CD19^+^ CD20^+^ IgD^−^ B cells were incubated with Total-seq C antibodies (Biolegend) for 30 min and washed 3 times. The number of cells were then counted and 14 000 cells per sample were run on the 10X Chromium (10X Genomics). Library preparation was completed by Biomedical Research Facility (BRF) at the JCSMR following the recommended protocols for the Chromium Single Cell 5′ Reagent Kit as well as 5’ Feature Barcode and V(D)J Enrichment Kit for Human B cells. Libraries were sequenced using the Illumina NovaSeq6000 (Illumina). The 10X Cell Ranger package (v1.2.0, 10X Genomics) was used to process transcript, CITE-seq and VDJ libraries and prepare them for downstream analysis. Details of all key reagents for single cell RNA-seq are given in Table S3.

## Quantification and statistics

### Single Cell RNA-seq analysis

The package *Seurat* (version 3.1) (Butler et al., 2018) was used for graph-based clustering and visualizations. All functions described are from *Seurat* or the standard R package (version 3.60) using the default parameters unless otherwise stated. Each sample was initially analyzed separately using the following procedures. Cells that expressed less than 200 genes and genes that were expressed in less than 3 cells were excluded, along with cells that had greater than 10% mitochondrial genes. Gene expression was normalized for both mRNA and CITE-seq assays using the NormalizeData function, then the 2000 most variable genes for each sample were identified using FindVaraibleFeatures. Next expression of all genes was scaled using ScaleData to linearly regress out sources of variation. Principal component analysis on the variable genes identified above was then run with RunPCA. Based on ElbowPlot results we decided to use 13, 20, 12 and 20 principal components (PCs) for the clustering of samples Non-Exp 1, Non-Exp 2, Exp 1 and Exp 2 respectively using FindNeighbours. FindClusters was then run to identify clusters for each sample, using the resolutions .3, .4, .5 and .4 respectively. FindAllMarkers was then used to identify clusters of non-B cells. The remaining cells in each sample were then normalized and scaled again as above. Australian and Kenyan samples were combined together first using FindIntegrationAnchors and then Intergratedata to create two combined datasets, 1 with both non-exposed samples and one with both malaria-exposed samples. These two combined samples were further combined using the commands to from one combined dataset containing all 4 samples. The combined dataset was then scaled again as above and a PCA was run. Using FindClusters with a resolution of 0.8, we identified our 11 clusters. DEGs were identified using FindAllMarkers. The clustering was visualized with Uniform Manifold Approximation and Projection (UMAP) dimensionality reduction using RunUMAP and plotted using DimPlot with umap as the reduction. Phylogentic analysis was done using BuildClusterTree to report the hierarchical distance matrix relating an ‘average’ cell from each cluster. Log-normalized gene expression data was visualized using violin plots (VlnPlots) as well as onto -UMAP plots (FeaturePlot). Heatmaps were generated using DoHeatmap.

For Smart-seq2 analysis, cells with greater than 10% mitochondrial genes were not excluded. 8 PCs were used as determine by ElbowPlot. For clustering a resolution of 0.8 was used.

### Diffusion Map Analysis

To create the diffusion map we utilized the R package *destiny* (Angerer et al., 2016). Our Seurat object was converted into a SingleCellExperiment object using as.SingleCellExperiment. The diffusion map was then generated using *Destiny’s* DiffusionMap command.

### Pseudotime Analysis

Pseudotime analysis was preformed using the R package *Monocle 3*. Our Seurat Object was converted into monocle3 main data calls cell_data_set. The default *Monocle 3* workflow was then followed.

### Gene Set Enrichment Analysis (GSEA)

GSEA was done using javaGSEA through the Broad Institute. For each comparison, DEGs were ranked by log-fold change and pre-ranked analysis using 1000 permutations was used to examine enrichment in selected gene sets (SUPP table?).

### VDJ Analysis

To determine the antigen-specific BCR repertoire, we made use of VDJpuzzle (Rizzetto et al., 2018) to reconstruct full-length heavy and light chains from each cell from out Smart-seq2 dataset. From this we were able to determine V region usage and mutation frequency.

VDJ sequences from the 10x dataset were obtained using the cellranger vdj command. From this output, V region usage and mutation frequency could be determined.

### Statistical Analysis

Statistical analysis of flow cytometry data was performed in GraphPad Prism for simple analyses without blocking factors; all other analyses was performed in R (The R Foundation for Statistical Computing) with details of statistical tests in the relevant figure legends. Abbreviations for p values are as follows: p < 0.05 = *, p < 0.01 = **, p < 0.001 = ***, p < 0.0001 = ****; with only significant p values shown.

### Data Deposition

Single cell RNA-seq data are deposited at NCBI BioProject accession number PRJNA612353: https://dataview.ncbi.nlm.nih.gov/object/PRJNA612353?reviewer=bf7ee45b186vstua0d23qud1nk

