## Supplementary Information for "Atypical B cells are a normal component of immune responses to vaccination and infection in humans"

**This Supplementary Information Includes**

Figures S1-S6

Tables S1-S3

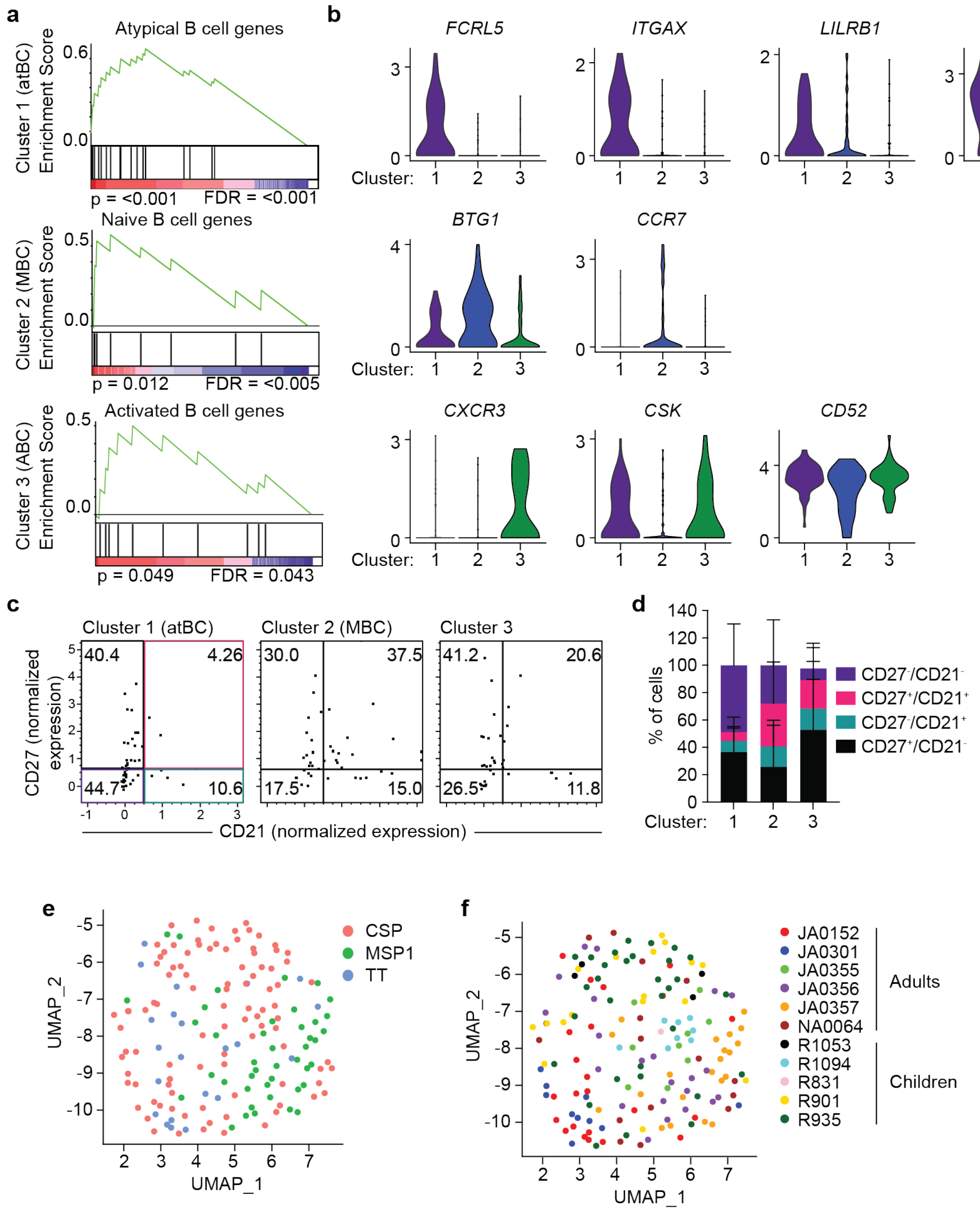


**Supplementary Figure 1: Tripartite division of antigen-specific B cells (related to Figure 1) A.** Representative GSEA plots of single cell RNA-seq data. Each graph shows genes associated with either atypical, naïve or activated B cells are enriched in either cluster 1,2 or 3 relative to all other cells **B.** Violin plots of the log-normalized expression of genes associated with atypical, classical or activated B cells by cluster, highest and lowest expression level noted. Each point represents a single cell and is colored by antigen-specificity. **C.** Scatter plots showing the MFI of CD27 and CD21 from index sort information including all cells analyzed, normalized across experiments. **D.** quantification of data shown in (C) separated by individual sample, bars are the mean % ± s.d.. **E.** Transcriptomes of antigen-specific B cells identified by Smart-seq2 methodology and visualized using UMAP. **F.** Transcriptomes of antigen-specific B cells identified by Smart-seq2 methodology and visualized using UMAP. Each point represents a single cell and is colored by sample.

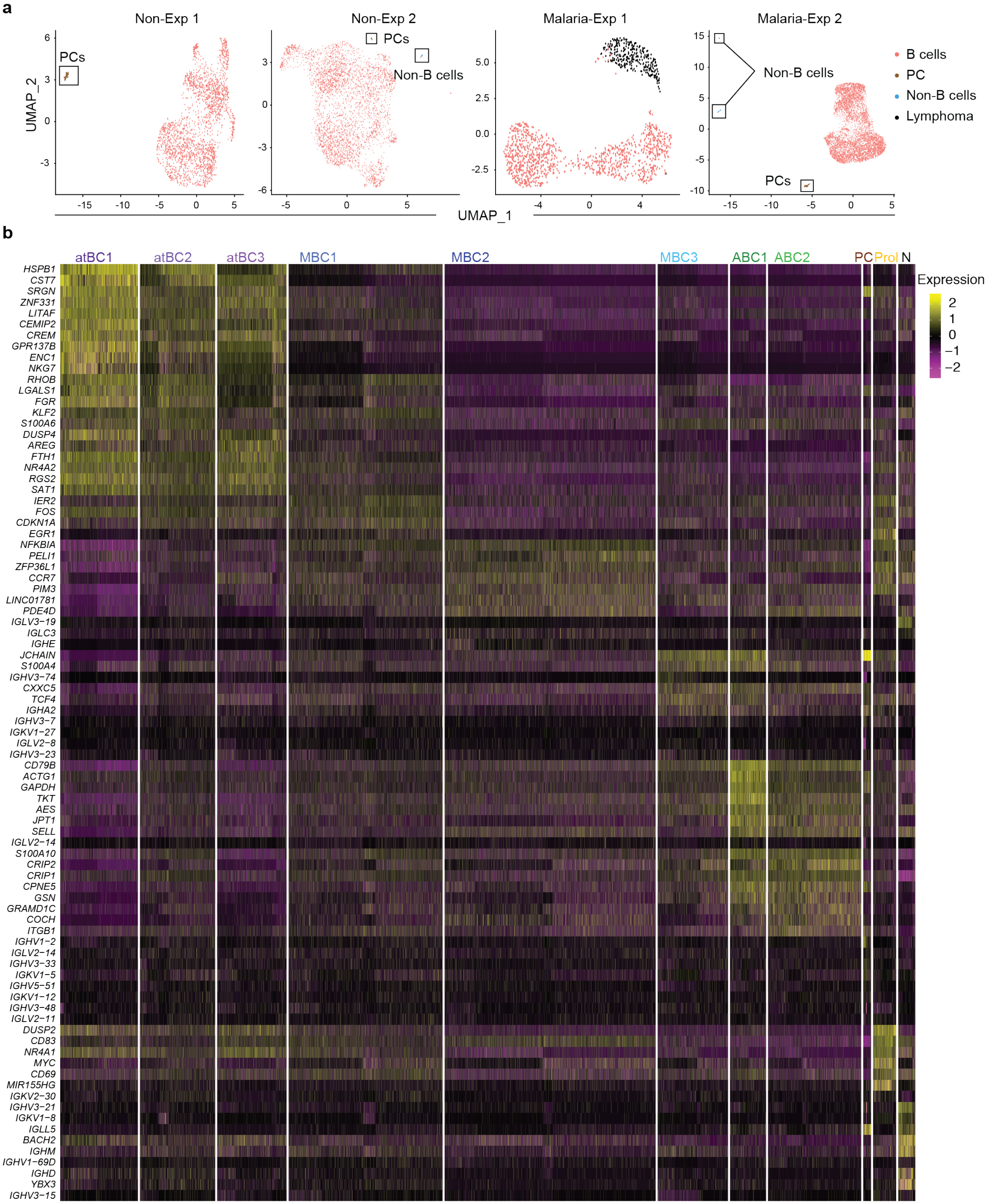


**Figure S2: Top DEGs per cluster identified by 10x Chromium single cell RNA-seq of circulating B cells (related to Figure 2).** Single B cells were sorted from 2 malaria exposed Kenyans individuals and 2 Australian individuals and gene expression was assessed using 10x chromium methodology. **A.**

Unsupervised clustering of sequenced cells from each individual visualized using UMAP. Each cell is represented by a point and colored based on cell type. **B.** Heatmap showing the expression of the top 10 DEGs (row) per cluster for each cell (column).

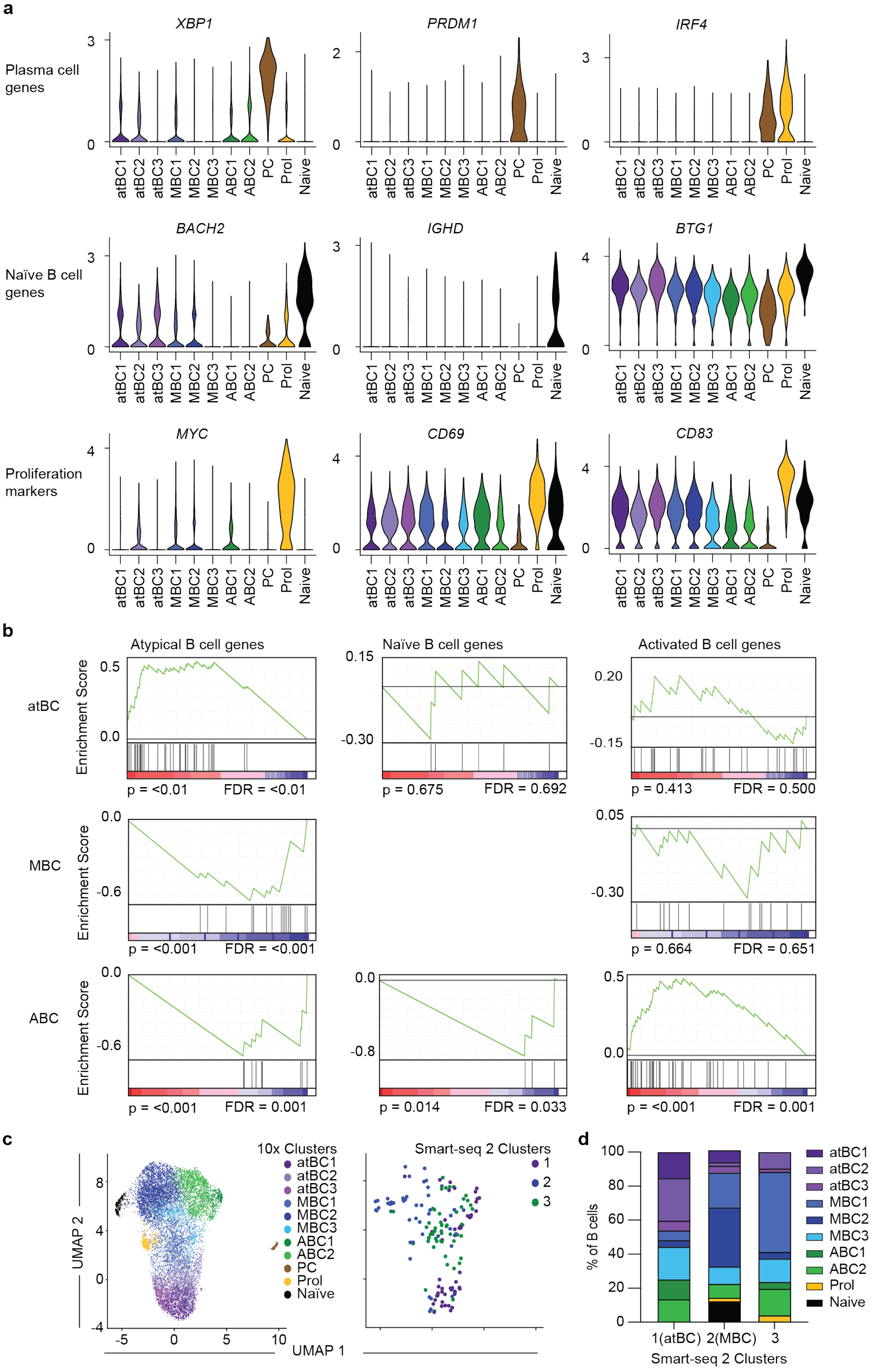
a
Plasma cell genes
XBP1
PRDM1
IRF4
Naïve B cell genes
BACH2
IGHD
BTG1
Proliferation markers
MYC
CD69
CD83
atBC1
atBC2
atBC3
MBC1
MBC2
MBC3
ABC1
ABC2
PC
Prol
Naive
b
Atypical B cell genes
Naïve B cell genes
Activated B cell genes
atBC
MBC
ABC
Enrichment Score
p = <0.01 FDR = <0.01
p = 0.675 FDR = 0.692
p = 0.413 FDR = 0.500
p = <0.001 FDR = <0.001
p = 0.664 FDR = 0.651
p = <0.001 FDR = 0.001
p = 0.014 FDR = 0.033
p = <0.001 FDR = 0.001
c
10x Clusters
Smart-seq 2 Clusters
UMAP 1
UMAP 2
Naïve
d
% of B cells
1(atBC)
2(MBC)
3
Smart-seq 2 Clusters

**Figure S3: Defining circulating B cell subsets from single cell RNA-seq data (related to Figure 2)** **A.** Violin plots of the log-normalized expression of genes associated with PCs, Naïve B cells or proliferating B cells by cluster, highest and lowest expression level noted. **B**. GSEA plots of single cell RNA-seq data. Each graph shows genes associated with either atypical, naïve or activated B cells that are enriched in either the atBC clusters, the MBC clusters or the ABC clusters relative to all other cells. **C.** Unsupervised clustering of B cells from an integrated 10x and Smart-seq 2 dataset visualized using UMAP split by sequencing technique. Each cell is represented by a point and colored by either 10x or Smart-seq2 clusters. **D.** Percentage of cells from each Smart-seq 2 cluster found in each 10x Chromium cluster.

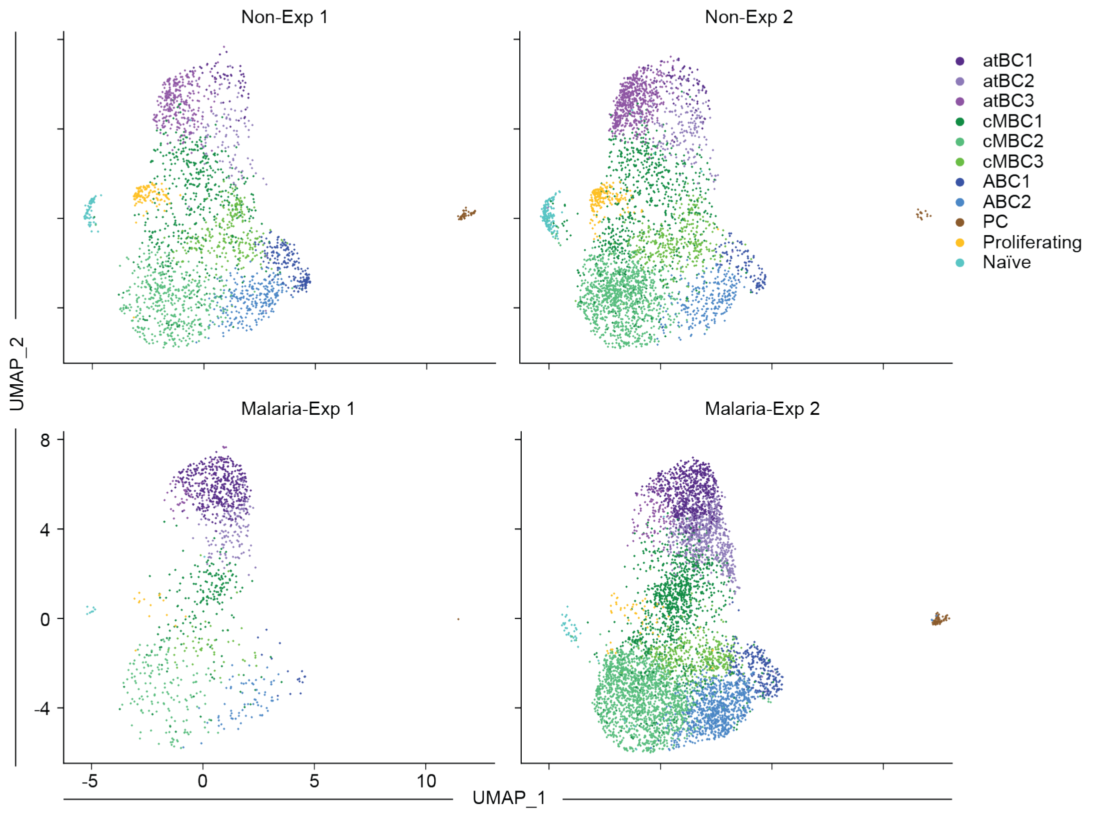


**Figure S4: Individual-level analysis of the number of cells per cluster (related to Figure 4) A.** Unsupervised clustering of circulating mature IgD$^{-}$ B cells visualized using UMAP, broken down by individual. Each cell is represented by a point and colored by cluster.

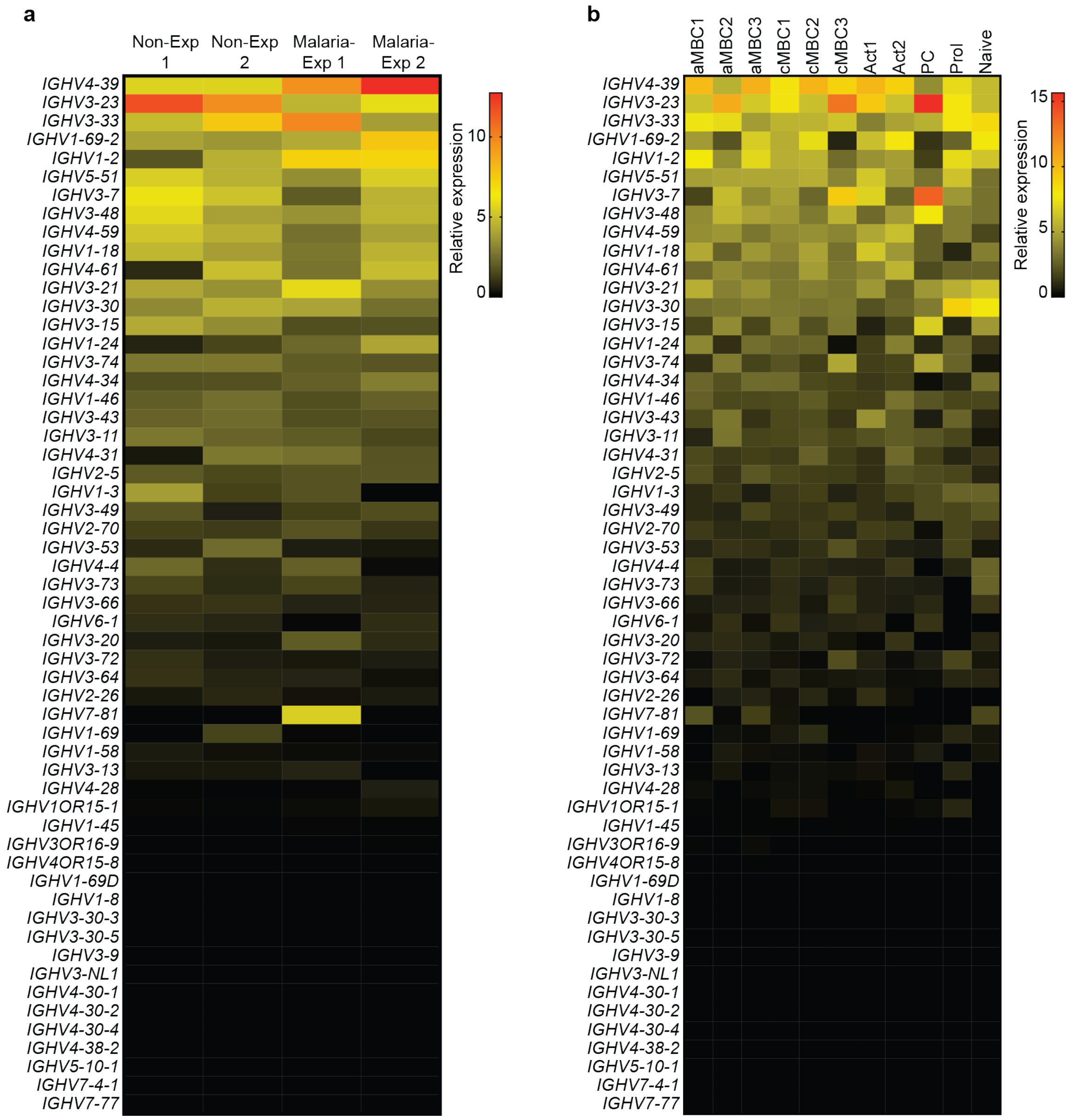


**Figure S5: *IGHV* gene usage among circulating B cells (related to Figure 5) A.** Heatmap of *IGHV* gene usage between individuals, ordered from the highest average expression to lowest. **B.** Heatmap of *IGHV* gene usage between clusters, ordered from the highest average expression to lowest.

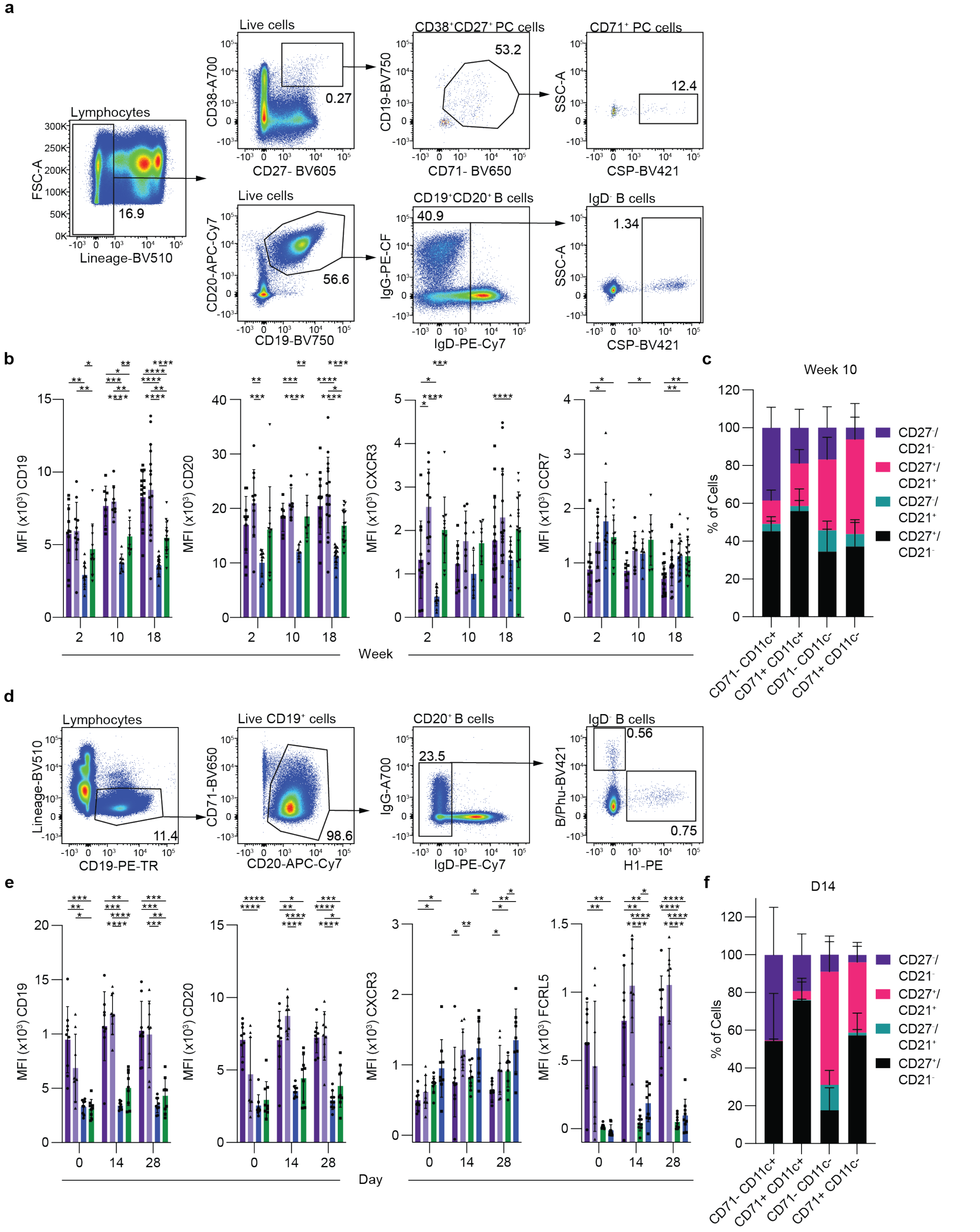

a
Lymphocytes
FSC-A
Lineage-BV510
16.9
Live cells
CD38-A700
CD27- BV605
0.27
CD38⁺CD27⁺ PC cells
CD19-BV750
CD71- BV650
53.2
CD71⁺ PC cells
SSC-A
CSP-BV421
12.4
Live cells
CD20-APC-Cy7
CD19-BV750
56.6
CD19⁺CD20⁺ B cells
IgG-PE-CF
IgD-PE-Cy7
40.9
IgD⁻ B cells
SSC-A
CSP-BV421
1.34
b
MFI (x10³) CD19
MFI (x10³) CD20
MFI (x10³) CXCR3
MFI (x10³) CCR7
Week
c
Week 10
% of Cells
CD27⁻/CD21⁻
CD27⁺/CD21⁺
CD27⁻/CD21⁺
CD27⁺/CD21⁻
CD71- CD11c+
CD71+ CD11c+
CD71- CD11c-
CD71+ CD11c-
d
Lymphocytes
Lineage-BV510
CD19-PE-TR
11.4
Live CD19⁺ cells
CD71-BV650
CD20-APC-Cy7
98.6
CD20⁺ B cells
IgG-A700
IgD-PE-Cy7
23.5
IgD⁻ B cells
B/Phu-BV421
H1-PE
0.56
0.75
e
MFI (x10³) CD19
MFI (x10³) CD20
MFI (x10³) CXCR3
MFI (x10³) FCRL5
Day
f
D14
% of Cells
CD27⁻/CD21⁻
CD27⁺/CD21⁺
CD27⁻/CD21⁺
CD27⁺/CD21⁻
CD71- CD11c+
CD71+ CD11c+
CD71- CD11c-
CD71+ CD11c-

**Figure S6: Characterizing the antigen specific B cell response following immunization (related to Figure 7) A.** Gating strategy to analyze CSP-specific B cells and PCs from PBMCs of donors immunized with PfSPZ vaccines. **B.** MFI of atypical and other markers found on CSP-specific B cells. **C.** Percentage of CSP-specific cells separated by expression of CD27 and CD21 on each cell type **D.** Gating strategy to analyze influenza-specific B cells from PBMCs of donors immunized with inactivated influenza vaccine. **E.** MFI of atypical and other markers found on influenza-specific B cells. **F.** Percentage of influenza-specific cells separated by expression of CD27 and CD21 on each cell type.

**Table S1: Demographic information for single cell RNA-seq donors**

| Sample | Age | Sex |
|---|---|---|
| JAO152 | 39 | Female |
| JAO164 (Exp 1) | 41 | Female |
| JAO301 | 42 | Male |
| JAO322 (Exp 2) | 42 | Male |
| JAO355 | 26 | Male |
| JAO356 | 25 | Male |
| JAO357 | 33 | Male |
| NA0064 | 30 | Female |
| R1053 | 2 | Male |
| R1094_6.5_months | 2 | Male |
| R1094_74_months | 8 | Male |
| R831 | 8 | Male |
| R901_6.5_months | 2 | Female |
| R901_74_months | 8 | Female |
| R935_6.5_months | 2 | Female |
| R935_74_months | 8 | Female |
| Non-Exp 1 | 26 | Female |
| Non-Exp 2 | 23 | Male |

**Table S2: Antibodies used in this study**

| Antibodies | Company | Catalog # |
|---|---|---|
| Anti-human CCR7 (Clone G043h7) BV650 | Biolegend | 353234 |
| Anti-human CD10 (Clone HI10a) BV421 | Biolegend | 312218 |
| Anti-human CD10 (Clone HI10a) BV510 | Biolegend | 312220 |
| Anti-human CD11c (Clone B-ly6) BUV737 | BD Biosciences | 741827 |
| Anti-human CD11c (Clone Bu15) BV421 | Biolegend | 337226 |
| Anti-human CD11c (Clone Bu15) PE | Biolegend | 337216 |
| Anti-human CD11c (Clone Bu15) PerCP Cy5.7 | Biolegend | 337216 |
| Anti-human CD14 (Clone M5E2) BV510 | Biolegend | 301842 |
| Anti-human CD14 (Clone M5E2) BV510 | Biolegend | 301842 |
| Anti-human CD14 (Clone M5E2) FITC | Biolegend | 301804 |
| Anti-human CD15 (SEA-1) (Clone H198) FITC | Biolegend | 394706 |
| Anti-human CD16 (Clone 3G8) BV510 | Biolegend | 302048 |
| Anti-human CD183 (CXCR3) (Clone G025H7) BV711 | Biolegend | 353732 |
| Anti-human CD184 (CXCR4) (Clone 12G5) BuV395 | BD Bioscience | 563924 |
| Anti-human CD19 (Clone HIB19) BV750 | Biolegend | 302261 |
| Anti-human CD19 (Clone J3-119) ECD | Beckman Coulter | IM2708U |
| Anti-human CD19 (Clone SJ25C1) BV605 | BD Bioscience | 562654 |
| Anti-human CD197 (CCR7) (Clone G043H7) BV785 | Biolegend | 353230 |
| Anti-human CD20 (Clone 2H7 ) APC Cy7 | BD Bioscience | 560853 |
| Anti-human CD20 (Clone 2H7) APC-Cy7 | BIolegend | 302314 |
| Anti-human CD21 (Clone B-ly4) BV711 | Biolegend | 563163 |
| Anti-human CD21 (Clone Bu32) PeCy7 | Biolegend | 354912 |
| Anti-human CD27 (Clone O323) BV605 | BioLegend | 302830 |
| Anti-human CD27 (Clone M-T271) PerCP Cy5.6 | Biolegend | 356408 |
| Anti-human CD3 (Clone HIT3A) FITC | Biolegend | 300306 |
| Anti-human CD3 (Clone OKT3) BV510 | Biolegend | 317332 |
| Anti-human CD307e (FCRL5) (Clone 509f6) APC | Biolegend | 340306 |
| Anti-human CD38 (Clone HIT2) Ax700 | BD Pharmingen | 560676 |
| Anti-human CD56 (Clone NCAM HCD56) BV510 | Biolegend | 318340 |
| Anti-human CD56 (NCAM) (Clone HCD56) FITC | Biolegend | 318304 |
| Anti-human CD71 (Clone CU1G4) BV650 | Biolegend | 334108 |
| Anti-human CD71 (Clone M-A712) FITC | BD Biosciences | 555536 |
| Anti-human CD8 (Clone RPA-T8) BV510 | Biolegend | 301048 |
| Anti-human CD83 (Clone Hb15e) PE | Biolegend | 305308 |
| Anti-human CXCR3 (Clone G025H7) APC | Biolegend | 353708 |
| Anti-human CXCR4 (Clone 12G5) BUV395 | BD Biosciences | 563924 |
| Anti-human IgD (Clone IA6-2) PE-CF594 | BD Bioscience | 562540 |

| | | |
|---|---|---|
| Anti-human IgD (Clone IA62) PE-Cy7 | BD Pharmingen | 561314 |
| Anti-human IgG Clone G18-145) PE-CF594 | BD Biosciences | 562538 |
| Anti-human IgG (Clone G18-145) A700 | BD Bioscience | 555784 |
| Anti-human IgM (Clone UCA-B1) BB700 | BD Pharmingen | 747879 |
| Anti-human Tbet (Clone 4B10) PerCP Cy5.5 | Biolegend | 644806 |

**Table S3: Key reagents used in this study**

| Reagents | Company | Identifer |
|---|---|---|
| Agencourt AMPure XP beads | Beckman Coulter | A63880 |
| Betaine 5M | Sigma-Aldrich | B300-1VL |
| CloneAmp HiFi PCR Premix | Takara | 639298 |
| ERCC Spike-In Mix | ThermoFisher | 4456740 |
| Nextera XT DNA Library Preparation kit | Illumina | FC-131-1096 |
| Nextera XT Index Kit v2 Set A (96 indices, 384 samples) | Illumina | FC-131-2001 |
| Nextera XT Index Kit v2 Set D (96 indices, 384 samples) | Illumina | FC-131-2004 |
| Quant-iT PicoGreen dsDNA Assay Kit | ThermoFisher | P11496 |
| RNAse inhibitor | Clontech | 2313A |
| SuperScript II RT | Life Technologies | 18064-071 |
| Triton X-100 | Sigma-Aldrich | T9284 |
| Chromium Next GEM chip G Single Cell Kit | 10x Genomics | 1000127 |
| Chromium Next GEM Single5' Library and Gel Bead Kit v1.1 | 10x Genomics | 1000080 |
| Chromium Single Cell 5' Feature Barcode Library Kit | 10x Genomics | 1000020 |
| Chromium Single Cell 5' Library Contruction Kit | 10x Genomics | 1000016 |
| Chromium Single Cell V(D)J Enrichment Kit Human B cell | 10x Genomics | 1000005 |
| Single Index Kit N Set A | 10x Genomics | 1000212 |
| Single Index Kit T Set A | 10x Genomics | 1000213 |
